# Analysis of combinatorial chemokine receptor expression dynamics using multi-receptor reporter mice

**DOI:** 10.1101/2021.08.11.455927

**Authors:** Laura Medina-Ruiz, Robin Bartolini, Douglas P Dyer, Francesca Vidler, Catherine E Hughes, Fabian Schuette, Samantha Love, Jun Fu, A. Francis Stewart, Gerard J Graham

## Abstract

Inflammatory chemokines and their receptors are central to the development of inflammatory/immune pathologies. The apparent complexity of this system, coupled with lack of appropriate in vivo models, has limited our understanding of how chemokines orchestrate inflammatory responses and has hampered attempts at targeting this system in inflammatory disease. Novel approaches are therefore needed to provide crucial biological, and therapeutic, insights into the chemokine-chemokine receptor family. Here, we report the generation of transgenic multi-chemokine receptor reporter mice in which spectrally-distinct fluorescent reporters mark expression of CCRs 1, 2, 3 and 5, key receptors for myeloid cell recruitment in inflammation. Analysis of these animals has allowed us to define, for the first time, individual and combinatorial receptor expression patterns on myeloid cells in resting and inflamed conditions. Our results demonstrate that chemokine receptor expression is highly specific, and more selective than previously anticipated.

## Introduction

Chemokines, and their receptors, are primary regulators of in vivo leukocyte migration and central orchestrators of innate and adaptive immune responses(Griffith, Sokol, & Luster, 2014; Rot & von Andrian, 2004). Chemokines are defined by a conserved cysteine motif and subdivided into CC, CXC, CX_3_C and XC subfamilies according to the specific configuration of this motif(Bachelerie et al., 2014; Rot & von Andrian, 2004). Chemokines signal through seven-transmembrane (7TM) spanning G protein-coupled receptors expressed by immune and inflammatory cells(Bachelerie et al., 2014; Rot & von Andrian, 2004), and these receptors are named according to the subfamily of chemokines with which they interact (CCR, CXCR, XCR and CX_3_CR).

Chemokines and their receptors can be functionally classified as either homeostatic or inflammatory according to the in vivo contexts in which they function(Mantovani, 1999; Zlotnik & Yoshie, 2000). Inflammatory chemokines and their receptors mediate leukocyte recruitment to inflamed, damaged or infected sites and are prominent contributors to a large number of autoimmune and inflammatory diseases(Viola & Luster, 2008). Chemokine receptors therefore represent important therapeutic targets(Proudfoot, Bonvin, & Power, 2015). However, to date, only two chemokine receptor antagonists have been approved for therapeutic use (Plerixafor, targeting CXCR4, and Maraviroc, targeting CCR5), and no antagonists have yet been approved for treating immune/inflammatory disease(Bachelerie et al., 2014). There are a number of explanations for this failure(Schall & Proudfoot, 2011), prominent amongst which is the fact that inflammatory chemokine and chemokine receptor biology is highly complex. For example, multiple chemokines are simultaneously expressed at inflamed sites and these interact in a complex, and poorly understood, manner with different inflammatory chemokine receptors(Bachelerie et al., 2014). To complicate matters further, reports in the literature indicate that distinct leukocyte subsets simultaneously express multiple chemokine receptors(Bachelerie et al., 2014; Griffith et al., 2014). As a result of this complexity it is currently difficult to precisely define in vivo roles for inflammatory chemokine receptors and we therefore lack a clear understanding of their integrated involvement in the orchestration of the inflammatory response.

The current study focuses on four inflammatory CC chemokine receptors (iCCRs)(Dyer et al., 2019): CCR1, CCR2, CCR3 and CCR5, which exemplify the ligand-receptor interaction complexity noted above. Collectively, these receptors direct non-neutrophilic myeloid cell trafficking at rest and during inflammation(Pease, 2011; Shi & Pamer, 2011) and are key players in inflammatory diseases. However, the resting and inflamed expression of these receptors on individual leukocyte populations has not been clearly defined. In addition, their individual roles in leukocyte mobilisation, recruitment and intra-tissue dynamics are still unclear. The use of knockout mice to study their function has been confused by partial phenotypes and conflicting results(Bennett et al., 2007; Humbles et al., 2002; Ma et al., 2002; Pope, Zimmermann, Stringer, Karow, & Rothenberg, 2005; Rottman et al., 2000; Tran, Kuziel, & Owens, 2000), leading to suggestions of redundancy in their function. There is therefore a pressing need to develop powerful novel tools to precisely define the temporo-spatial patterns of expression of these receptors at rest and during inflammation and in so doing to precisely delineate their roles in the physiological and pathological inflammatory response.

Here we report the generation of an iCCR reporter (iCCR-REP) mouse strain expressing spectrally-distinct fluorescent reporters for CCR1, CCR2, CCR3 and CCR5. We have used these mice to provide, for the first time, a comprehensive analysis of iCCR expression on bone marrow (BM), peripheral blood and tissue-resident myeloid cells at rest and during inflammatory responses. In contrast to published data, our analysis indicates selective receptor expression in individual cell types, in resting and acute inflammatory contexts and suggests little, if any, redundancy in function. We propose that these mice represent a transformational addition to the suite of mouse models available for analysing inflammatory chemokine receptor function *in vivo* and that they will be instrumental in helping to deconvolute the complexity of the chemokine-driven inflammatory response in a variety of pathological contexts.

## Results

### Generation of iCCR-reporter mice

The *iCcrs* are organised in a single 170 kb genomic cluster, which contains no other genes and which is highly conserved among mammals(Nomiyama, Osada, & Yoshie, 2011) and located on mouse chromosome 9. To produce the iCCR-reporter (iCCR-REP) mice, we generated a recombineered version of a bacterial artificial chromosome (BAC) encompassing the cluster (*iCcr*-REP cluster), in which the coding sequence of each of the *iCcr*s was replaced with sequences encoding spectrally-distinct fluorescent proteins. The reporters used in this study (mTagBFP2(Subach, Cranfill, Davidson, & Verkhusha, 2011), Clover, mRuby2(Lam et al., 2012) and iRFP682(Shcherbakova & Verkhusha, 2013)), were selected on the basis of their discrete excitation and emission spectra (Figure 1A). Using counterselection recombineering(Wang et al., 2014), *Ccr1* was replaced with Clover, *Ccr2* with mRuby2, *Ccr3* with mTagBFP2 and *Ccr5* with iRFP682 (Figure 1Bi and ii). Transgenic iCCR-REP mice were generated by pro-nuclear injection of the *iCcr*-REP BAC (Figure 1Biii). Using targeted locus amplification (TLA)(de Vree et al., 2014), the *iCcr*-REP cluster was located to chromosome 16:82389380-82392016 (Figure 1Ci) where 5-8 copies of the BAC were inserted in a head-tail manner. Insertion of the *iCcr*-REP clusters lead to the deletion of a 2.5 Kb genomic region that contained no coding or regulatory sequences (Figure 1Cii).

**Figure 1.**
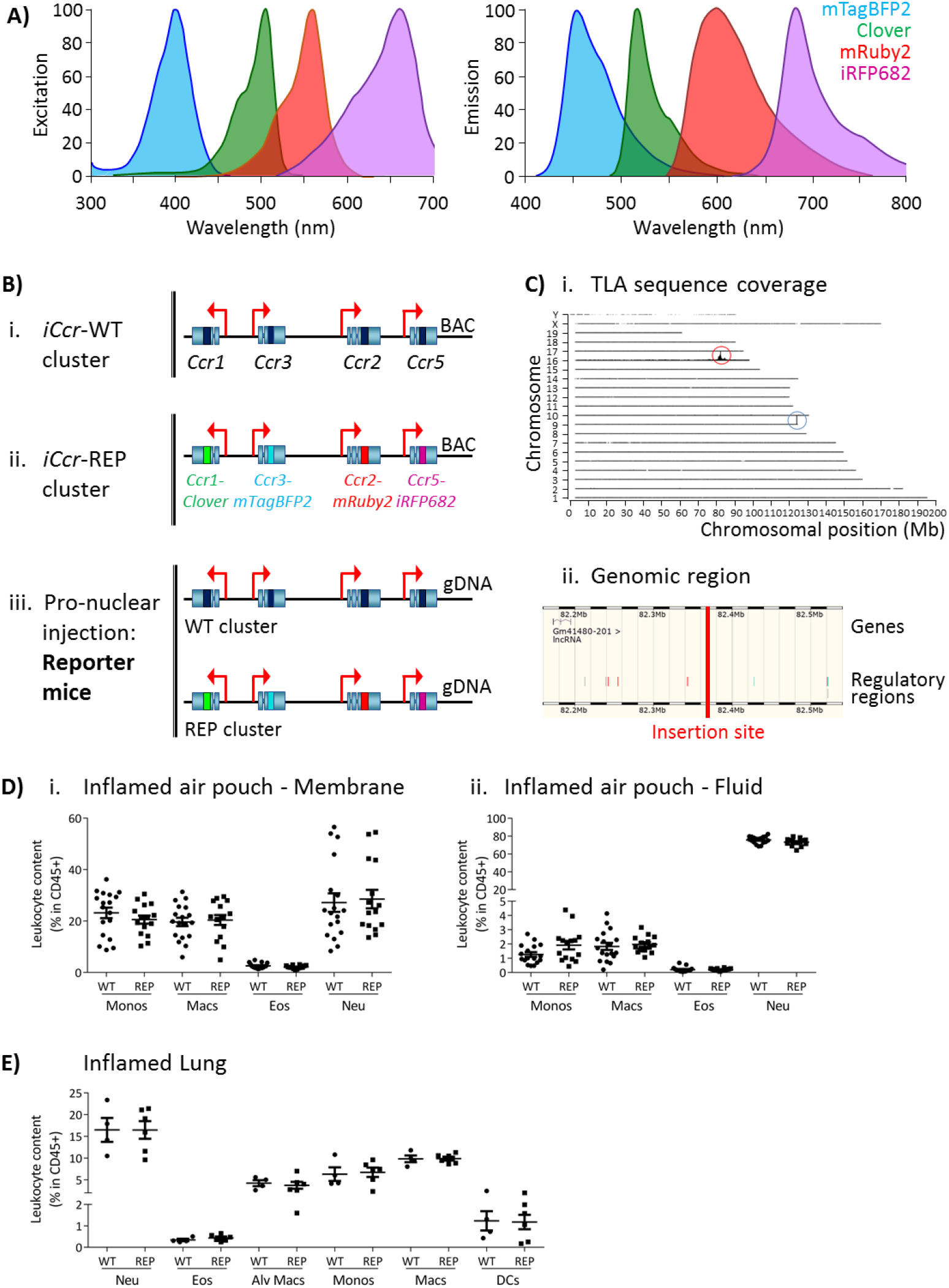
Generation of the reporter mice. A) Reporters were selected for this study based on their discrete excitation and emission spectra. B) (i) The *iCcrs* were targeted in a Bacterial Artificial Chromosome (BAC). (ii) The coding sequence of each *iCcr* was replaced with a different fluorescent reporter (ii). (iii) Pronuclear injection was then used to generate the transgenic reporter mice. C) (i) The *iCcr*-REP cluster inserted into chromosome 16 (red circle), as determined by targeted locus amplification (TLA). The blue circle represents the endogenous locus. (ii) The insertion site does not contain any other genes or regulatory regions. D) Leukocyte counts determined by flow cytometry in the inflamed air pouch. Data are shown for (i) the membrane that surrounds the air pouch and for (ii) the lavage fluid. E) Leukocyte counts determined by flow cytometry in inflamed lungs. Data on D and E are shown as mean ±SEM and are compiled from at least two separate experiments. Normally distributed data were analysed using unpaired t-test with or without Welch’s correction, according to their standard deviations. Not normally distributed data were analysed using Mann-Whitney or Kolmogorov-Smirnov, according to their standard deviations. Each data point represents a measurement from a single mouse. Abbreviations are: Monos, monocytes; Macs, macrophages; Neu, neutrophils; Eos, eosinophils; DCs, dendritic cells; Alv macs, alveolar macrophages. See also Figure S3.

Thus, using recombineering, we have generated transgenic (iCCR-REP) mice expressing spectrally-distinct reporters for each of the iCCRs.

The iCCR-REP mice maintain the original *iCcr* cluster on chromosome 9. To confirm that the transgene does not interfere with normal iCCR-dependent myeloid cell recruitment, we examined myeloid cell population sizes in different tissues by flow cytometry (gating strategies: Figures S1 and S2). Analysis of resting mice demonstrated that the myeloid cell content in these tissues was indistinguishable between iCCR-REP and wild type (WT) animals for all populations analysed (Figure S3). Next, we tested for possible effects of the transgene on myeloid cell recruitment to inflamed sites. To this end we used two different models. First, we used the air-pouch model of inflammation, involving the generation of an air cavity under the dorsal skin of the mouse and the injection of carrageenan into the cavity to induce inflammation. In line with the results obtained from resting tissues, analysis of the myeloid cell content in the membrane surrounding the inflamed air-pouch, as well as in the fluid collected from the inflamed cavity, showed no differences between iCCR-REP and WT animals (Figure 1D). We also analysed cellular content in inflamed lungs of mice that received an intranasal dose of lipopolysaccharide (LPS). Again, no differences were observed between iCCR-REP and WT animals in the size of any of the populations measured (Figure 1E).

Together, these data demonstrate that iCCR-REP mice have normal iCCR-dependent myeloid cell recruitment/migration dynamics at rest and under inflammatory conditions.

### iCCR-reporter expression accurately replicates iCCR surface presentation

To confirm that reporter expression faithfully recapitulates iCCR surface presentation, we compared iCCR antibody binding and iCCR reporter expression by flow cytometry (gating strategies: Figures S1A and S2B). As shown in Figure 2A, the majority of splenic Ly6C^hi^ monocytes from resting WT mice displayed anti-CCR2 antibody staining and essentially all mRuby2/CCR2 expressing Ly6C^hi^ monocytes in iCCR-REP mice were co-positive for CCR2 antibody staining. Background autofluorescence was undetectable in WT or iCCR-deficient (iCCR-def) mice(Dyer et al., 2019) although we routinely detected non-specific antibody staining on iCCR-def cells. Similar results were obtained when analysing splenic SiglecF^+^ eosinophils and kidney CD11b^+^F480^+^ macrophages for expression of mTagBFP2/CCR3 and iRFP682/CCR5 respectively. Splenic iCCR-REP eosinophils expressing mTagBFP2 simultaneously displayed surface CCR3 antibody staining (Figure 2B) and kidney iCCR-REP macrophages expressing iRFP682 simultaneously showed surface CCR5 antibody staining (Figure 2C). In both cases, low level non-specific antibody staining was observed on iCCR-def cells. Clover/CCR1 was detected on kidney CD11b^+^F480^+^ macrophages of iCCR-REP mice (Figure 2Di and ii). However, in our hands, no commercially available antibodies were able to detect CCR1 on any tissues analysed. For that reason, we used RNAscope to confirm expression of CCR1 on kidney cells. As shown in Figure 2Diii and iv, CCR1 transcripts were clearly detectable on WT and iCCR-REP, but not iCCR-def kidney sections.

**Figure 2.**
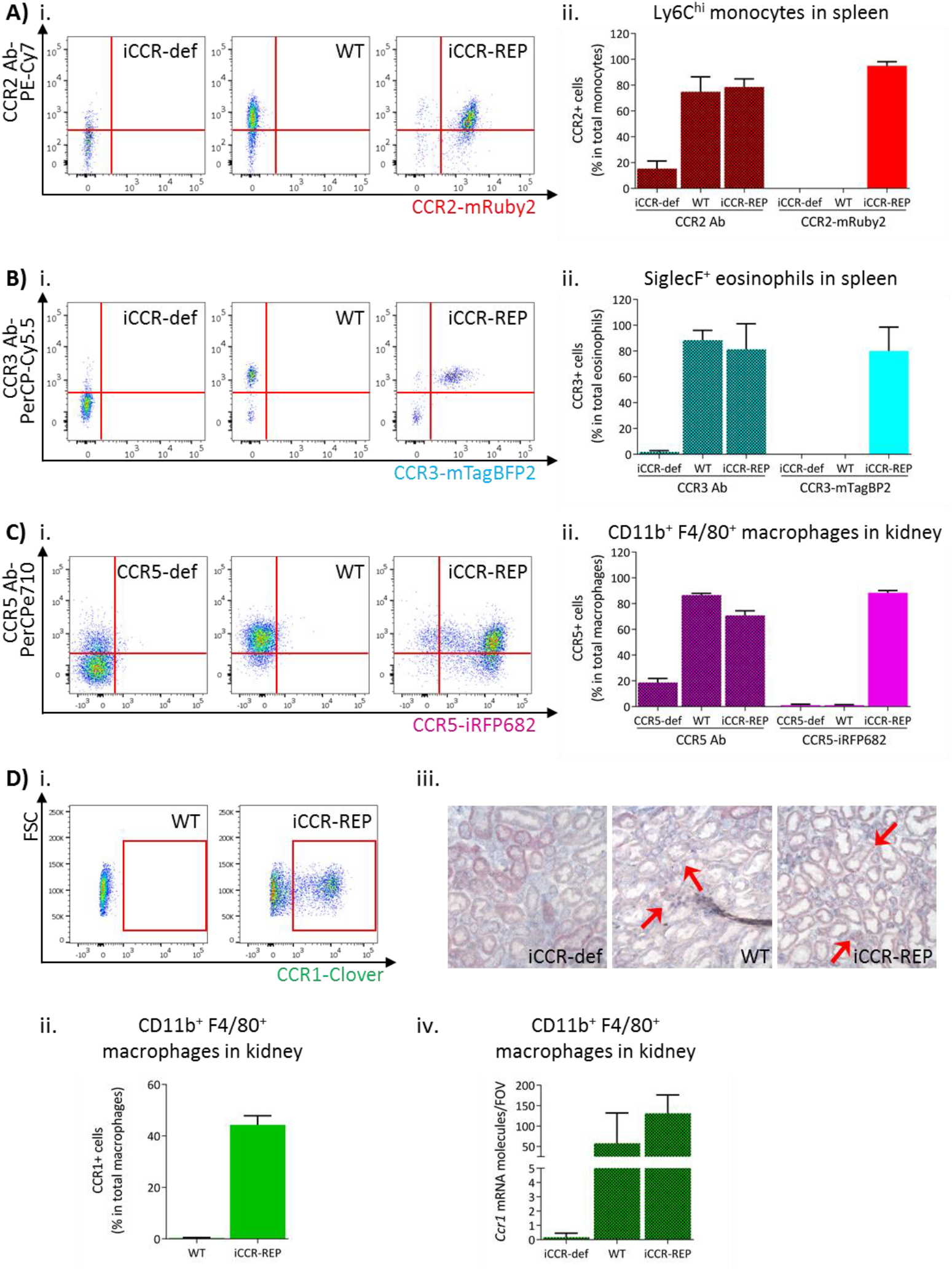
iCCR reporter expression accurately mirrors iCCR surface presentation. A) (i) Flow cytometric analysis of CCR2 antibody binding and mRuby2 expression in spleen Ly6C^hi^ monocytes at rest. (ii) Quantification of the percentage of Ly6C^hi^ monocytes binding CCR2 antibody and expressing mRuby2. B) (i) Flow cytometric analysis of CCR3 antibody binding and mTagBFP2 expression in spleen SiglecF^+^ eosinophils at rest. (ii) Quantification of the percentage of SiglecF^+^ eosinophils binding CCR3 antibody and expressing mTagBFP2. C) (i) Flow cytometric analysis of CCR5 antibody binding and iRFP682 expression in kidney CD11b^+^F4/80^+^ macrophages at rest. (ii) Quantification of the percentage of CD11b^+^F4/80^+^ macrophages binding CCR5 antibody and expressing iRFP682. D) (i) Flow cytometric analysis of Clover expression in kidney CD11b^+^F4/80^+^ macrophages at rest. (ii) Quantification of the percentage of CD11b^+^F4/80^+^ macrophages expressing Clover. (iii) Brightfield images of resting kidneys showing CCR1 mRNA molecules detected by RNAscope analysis in the form of red precipitate dots (arrows). (iv) CCR1 mRNA molecule counts per field of view. Data information: data on Aii, Bii, Cii, Dii and Div are shown as mean ±SD and are compiled from at least two separate experiments.

Thus, these data confirm that expression of the reporters in the iCCR-REP mice faithfully reflects iCCR expression and surface presentation. These mice, therefore, represent a unique, and validated, resource to examine individual and combinatorial iCCR expression in leukocytes.

### BM and circulating leukocytes express specific iCCRs

Next, we used flow cytometric analysis to examine iCCR expression in the iCCR-REP mice with WT littermates as controls for background autofluorescence. We first determined iCCR expression in myeloid cells from resting BM and blood. Cell suspensions were prepared from both compartments and stained with leukocyte subset-specific antibodies (gating strategies: Figure S1B). We initially assessed iCCR-reporter expression in Ly6C^hi^ monocytes. As expected, and in agreement with previous reports(Geissmann, Jung, & Littman, 2003; Saederup et al., 2010), the majority of Ly6C^hi^ monocytes were positive for mRuby2/CCR2, in BM and in blood. mTagBFP2/CCR3 and iRFP682/CCR5 were not detected on monocytes. However, Clover/CCR1 was seen on a small number of Ly6C^hi^ monocytes, representing approximately 3% of the population (Figure 3Ai-iv). Interestingly, in all cases, Clover was co-expressed with mRuby2 (Figure 3Av-vii).

**Figure 3.**
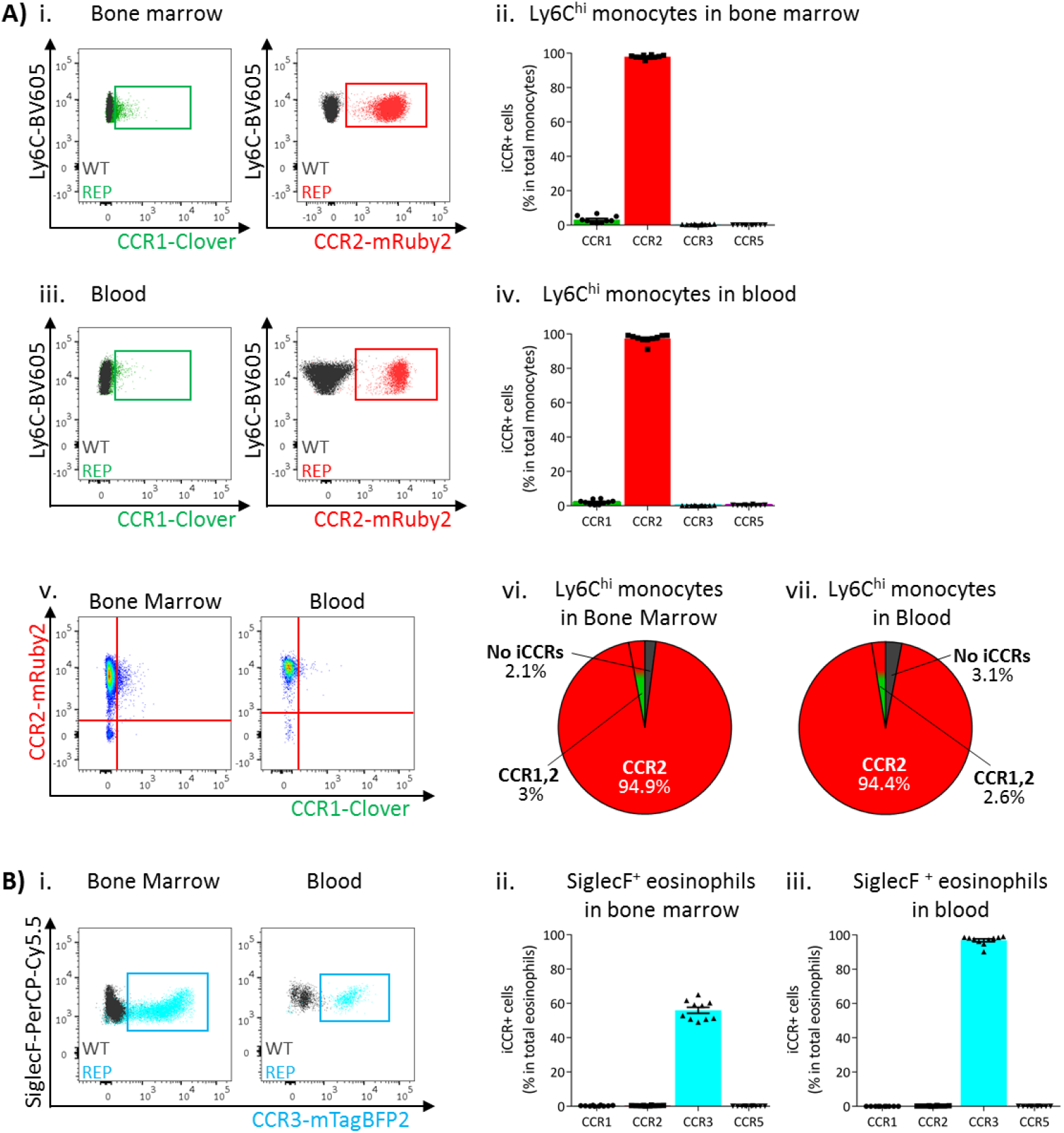
iCCR expression in bone marrow and blood leukocytes at rest. A) Flow cytometric analysis of Clover/CCR1 and mRuby2/CCR2 expression in (i) bone marrow and (iii) circulating Ly6C^hi^ monocytes. Quantification of the percentage of Ly6C^hi^ monocytes expressing the fluorescent reporters in (ii) bone marrow and (iv) blood. (v) Flow cytometric analysis of Clover/CCR1 and mRuby2/CCR2 co-expression in bone marrow and circulating Ly6C^hi^ monocytes. Distribution of Clover and mRuby2 in (vi) bone marrow and (vii) circulating Ly6C^hi^ monocytes. B) (i) Flow cytometric analysis of mTagBFP2/CCR3 expression in bone marrow and circulating SiglecF^+^ eosinophils. Quantification of the percentage of SiglecF^+^ eosinophils expressing the iCCR reporters in (ii) bone marrow and (iii) blood. Data on A–B are compiled from at least three separate experiments. Data on Aii, Aiv, Bii and Biii are shown as mean ±SEM (N=10). Each data point represents a measurement from a single mouse. Blots in Ai, Aiii and Bi are combinatorial blots showing reporter expression in iCCR REP and WT (control for background autofluorescence) mice.

SiglecF^+^ eosinophils exclusively expressed mTagBFP2/CCR3 (Figure 3Bi) from the reporter cluster. However, only approximately 50% of the population expressed this reporter in the BM (Figure 3Bii), while levels increased as cells entered the circulation, where the majority of eosinophils now expressed mTagBFP2 (Figure 3Biii). We did not detect expression of any of the iCCR reporters in resting Ly6G^+^ neutrophils (data not shown).

Thus, the iCCR-REP mice demonstrate that, with the exception of a small population of monocytic cells, individual iCCRs display selective association with discrete myeloid lineages in BM and peripheral blood.

### iCCR expression in resting tissues

Expression of the iCCRs was next assessed in resident myeloid cell populations of resting lungs and kidneys (gating strategies: Figure S2). The majority of Ly6C^hi^ monocytes from resting lungs expressed mRuby2/CCR2, with only a very small fraction co-expressing Clover/CCR1 or iRFP682/CCR5 (Figure 4Ai-iii). In contrast, a high proportion of the monocyte-derived CD11b^+^F480^+^MHCII^Lo^ interstitial macrophages (IMs) (Chakarov et al., 2019; Gibbings et al., 2017) expressed Clover/CCR1 and iRFP682/CCR5, with only a small fraction expressing mRuby2/CCR2 (Figure 4Bi-iv). Interestingly, in this case, CCR1 and CCR5 are mainly expressed independently of CCR2 (Figure 4Bv), suggesting that macrophages down-regulate CCR2 and induce CCR1 and CCR5 as they differentiate from infiltrating monocytes. In line with these findings, we observed that the mean fluorescence intensity (MFI) of mRuby2 in CCR2^+^ interstitial macrophages was lower than that in CCR2^+^ monocytes (Figure 4Bvi). This confirms that the CCR2 downregulation does not simply reflect reduction in the number of macrophages expressing the receptor but also reduced expression.

**Figure 4.**
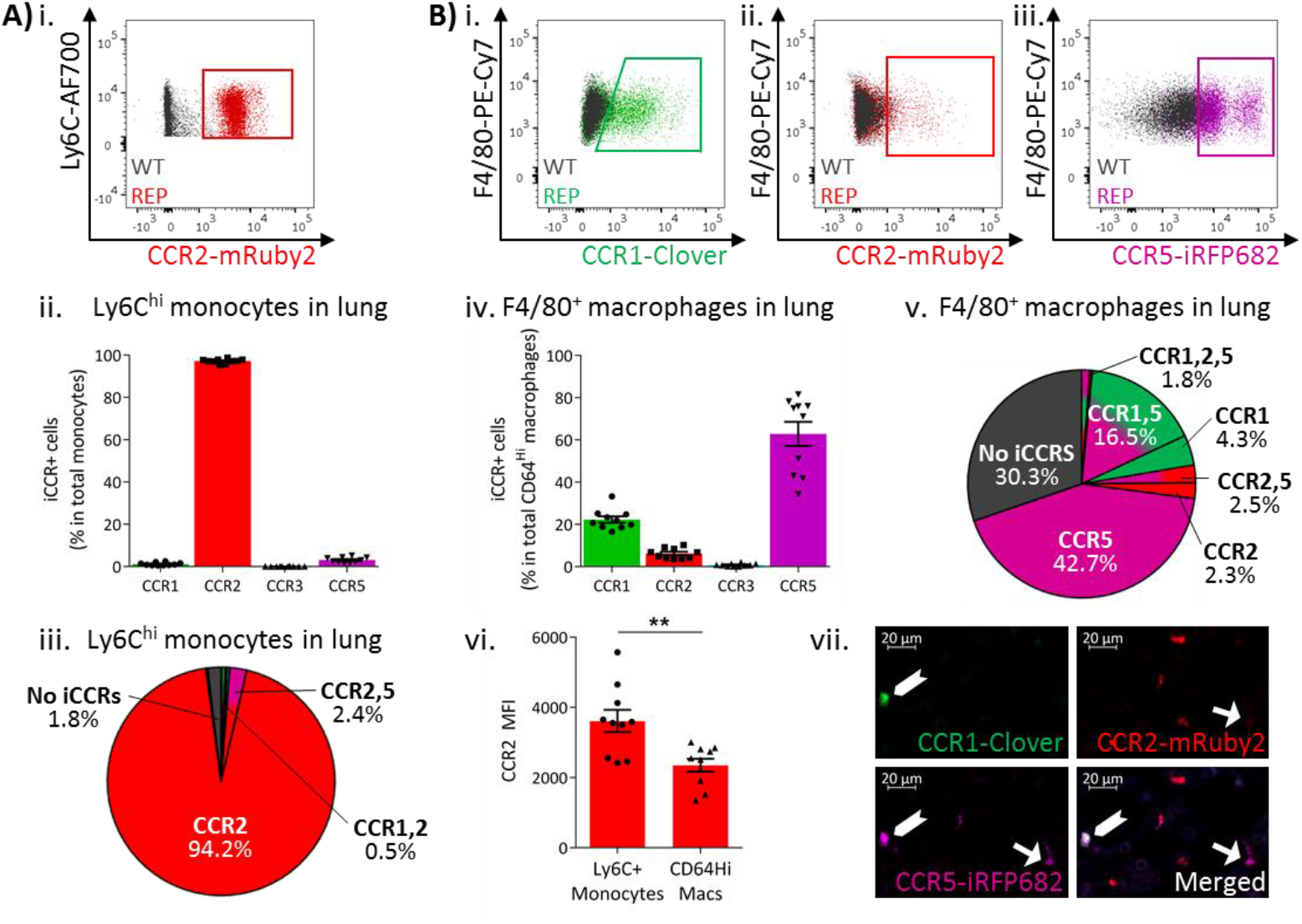
iCCR expression in resting lung. A) (i) Flow cytometric analysis of mRuby2/CCR2 expression in Ly6C^hi^ monocytes. (ii) Quantification of the percentage of Ly6C^hi^ monocytes expressing the iCCR reporters. (iii) Distribution of the iCCR reporters in Ly6C^hi^ monocytes. B) Flow cytometric analysis of (i) Clover/CCR1, (ii) mRuby2/CCR2 and (iii) iRFP682/CCR5 expression in F4/80^+^ macrophages. (iv) Quantification of the percentage of F4/80^+^ macrophages expressing the iCCR reporters. (v) Distribution of the iCCR reporters in F4/80^+^ macrophages. (vi) mRuby2/CCR2 mean fluorescence intensity from Ly6C^hi^ monocytes and F4/80^+^ macrophages. (vii) Lung leukocytes expressing CCR2 exclusively, CCR1 and CCR5 (chevron) or CCR2 and CCR5 (arrow). Data in A–B are compiled from at least two separate experiments. Data on Aii, Biv, Bvi are shown as mean ±SEM (N=10). Each data point represents a measurement from a single mouse. Blots in Ai, Bi, Bii and Biii are combinatorial blots showing reporter expression in iCCR REP and WT (control for background autofluorescence) mice. Data on Bvi were analysed using unpaired t-test. **p <0.01. See also Figure S4 and S5.

Similar results were obtained from resting kidneys. Again, Ly6C^hi^ monocytes expressed almost exclusively CCR2, with only some co-expressing CCR1 or CCR5 (Figure 5Ai-iii). However, monocyte-derived CD11b^+^F480^+^ macrophages (Puranik et al., 2018) mainly expressed CCR1 and CCR5, with only a small fraction retaining CCR2 expression (Figure 5Bi-v). Consistent with the observations in lung, the CCR2^+^ macrophages expressed lower CCR2 levels than Ly6C^hi^ monocytes as confirmed by the lower MFI for mRuby2 in this population (Figure 5Bvi). Together, these results suggest that this iCCR expression pattern is consistent across different tissues under resting conditions.

**Figure 5.**
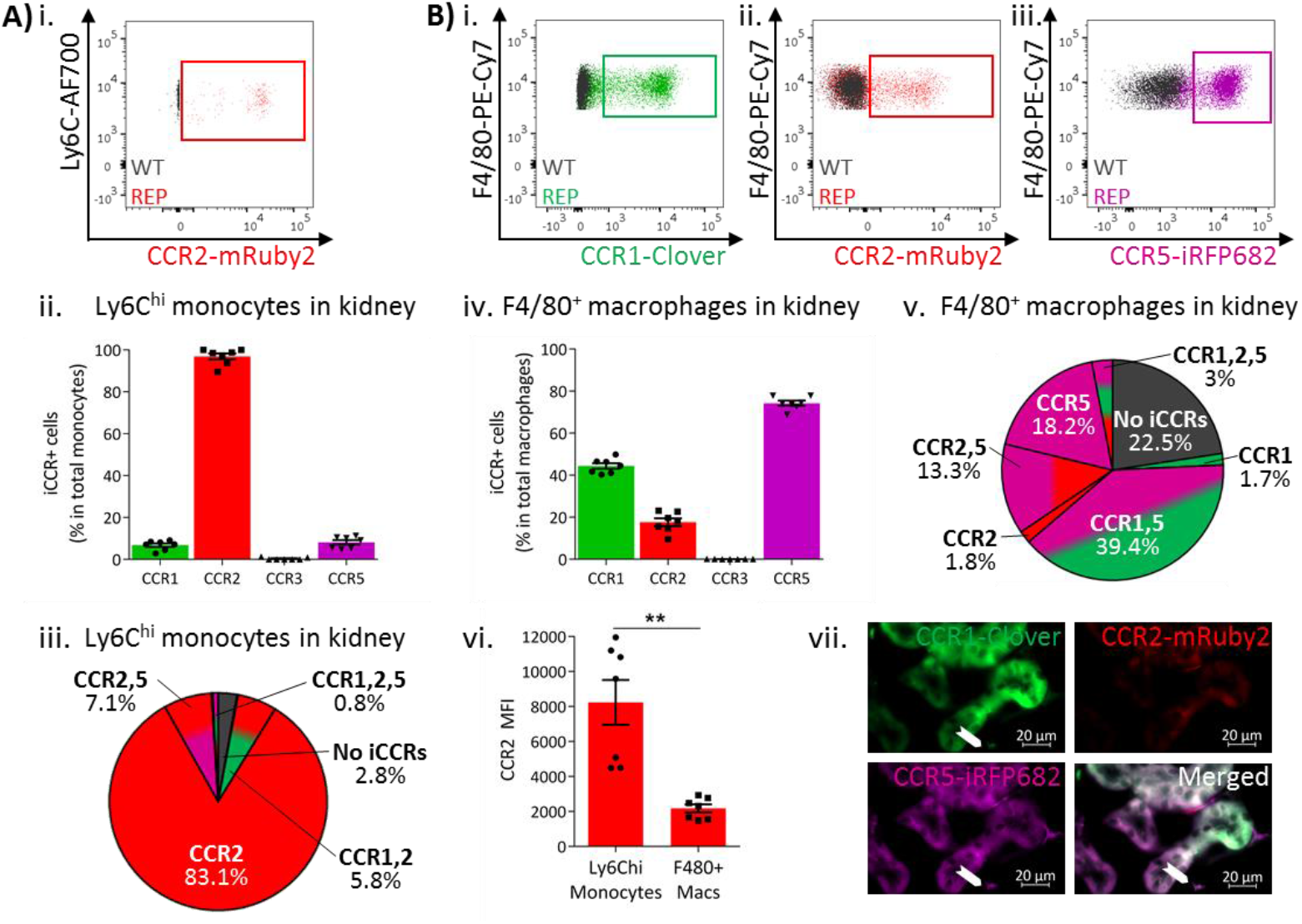
iCCR expression in resting kidney. A) (i) Flow cytometric analysis of mRuby2/CCR2 expression in Ly6C^hi^ monocytes. (ii) Quantification of the percentage of Ly6C^hi^ monocytes expressing the iCCR reporters. (iii) Distribution of the iCCR reporters in Ly6C^hi^ monocytes. B) Flow cytometric analysis of (i) Clover/CCR1, (ii) mRuby2/CCR2 and (iii) iRFP682/CCR5 expression on F4/80^+^ macrophages. (iv) Quantification of the percentage of F4/80^+^ macrophages expressing the iCCR reporters. (v) Distribution of the iCCR reporters in F4/80^+^ macrophages. (vi) mRuby2/CCR2 mean fluorescence intensity from Ly6C^hi^ monocytes and F4/80^+^ macrophages. (vii) Kidney leukocytes expressing CCR2 exclusively, CCR5 exclusively or CCR1 and CCR5 (chevron). Data in A–B are compiled from at least two separate experiments. Data on Cii, Civ and Cvi are shown as mean ±SEM (N=7). Each data point represents a measurement from a single mouse. Blots in Ai, Bi, Bii and Biii are combinatorial blots showing reporter expression in iCCR REP and WT (control for background autofluorescence) mice. Data on Bvi were analysed using unpaired t-test with Welch’s correction. **p <0.01. See also Figure S5.

We next assessed iCCR expression in lung CD11b^+^F480^+^MHCII^Hi^ IMs (Chakarov et al., 2019). As shown in Figure S4, F480^Lo^MHCII^Hi^ IMs expressed CCRs 1, 2 and 5 (Figure S4Ai-iv). F480^+^MHCII^Hi^ IMs expressed CCR5, but CCR1^+^ and CCR2^+^ cells were less abundant (Figure S4Bi-iv). This pattern is similar to that of MHCII^Lo^ IMs, suggesting that F480^+^MHCII^Hi^ IMs also downregulate CCR2 as they differentiate from F480^Lo^MHCII^Hi^ IMs. In line with this, co-expression of CCR1 or CCR5 with CCR2 was more apparent in F480^Lo^MHCII^Hi^ than in F480MHCII^Hi^ ^+^ IMs (Figure S4Ci-iv).

To further confirm this pattern, we generated GM-CSF derived macrophages from BM of iCCR-REP mice. As shown in Figure S4, and in line with previous findings, freshly extracted BM Ly6C^hi^ monocytes expressed CCR2 almost exclusively, with only a small fraction co-expressing CCR1 (Figure S4D). After 2 days in culture with GM-CSF, Ly6C^hi^ monocytes retain expression of CCR2 but induce CCR1 and CCR5, with approximately 40% of the cells co-expressing all three receptors (Figure S4D). At this time point, CD11c^+^ macrophage precursors are already detectable in cultures. These precursors express CCR1 and CCR5 to a higher extent than observed in Ly6C^hi^ monocytes, with 65% of the population co-expressing CCRs 1, 2 and 5 (Figure S4D). By day 9 in culture, fully mature macrophages express mainly CCRs 1 and 5, which are now detected independently from CCR2 in over 50% of the cells (Figure S4D).

At rest, lung eosinophils only expressed mTagBFP2/CCR3 (Figure S4Di-ii), whereas alveolar macrophages did not express any of the iCCR reporters (data not shown). We did not detect expression of any of the iCCR reporters in lung or kidney neutrophils (data not shown).

Thus, these data demonstrate that, whilst monocytes predominantly express CCR2, they downregulate this receptor and upregulate CCR1 and CCR5 as they differentiate. In keeping with the observations from blood and BM, eosinophils within tissues are positive only for CCR3.

### iCCR expressing cells can be directly visualised in tissues

The above analyses focused on flow cytometric evaluation of iCCR-REP expression. However, we next determined whether these mice are also appropriate for direct visualisation and localisation of iCCR-expressing cells within tissues. As shown in Figure S5, fluorescence imaging of a section from resting spleen revealed easily identifiable cells expressing each of the four reporters. In addition, combinatorial iCCR expression was detected on individual cells as shown in Figures 4Bvii. Here we highlight lung cells expressing only mRuby2/CCR2 as well as cells that are co-expressing Clover/CCR1 with iRFP682/CCR5 or mRuby2/CCR2 with iRFP682/CCR5. Notably, the cells expressing exclusively CCR2 are brighter for mRuby2 than the CCR2^+^ CCR5^+^ cells, consistent with results above suggesting that macrophages downregulate CCR2 expression as they upregulate CCR1 or CCR5. Finally, the imaging approaches are applicable to numerous tissues and, as shown in Figure 5Bvii, iCCR-REP mice can be used to identify leukocyte populations expressing individual and combinatorial patterns of iCCR expression in the resting kidney.

### iCCR expression in BM and circulation under inflammatory conditions

To determine myeloid cell iCCR expression under inflamed conditions we first used the air-pouch model (Figure 6A). iCCR expression in BM and blood was assessed 48 hours after the induction of inflammation (gating strategies: Figure S1B). Similar to resting conditions, the majority of Ly6C^hi^ monocytes expressed mRuby2/CCR2 and a fraction expressed Clover/CCR1 (Figure 6Bi-ii and 6Biv-v). However, the fraction of Ly6C^hi^ monocytes expressing CCR1 was higher than in resting mice. Approximately 20% of Ly6C^hi^ monocytes expressed CCR1 in the BM of inflamed mice, representing a 6-fold increase compared to resting conditions (Figure 6Biii). Similarly, in blood, approximately 11% of Ly6C^hi^ monocytes expressed CCR1, representing a 5-fold increase compared with resting conditions (Figure 6Bvi). Interestingly, in this inflamed context, we also detected, for the first time, expression of CCR1 independently of CCR2 on monocytes, although in most cases both iCCRs are co-expressed (Figure 6Ci-iii). BM and blood eosinophils showed similar iCCR expression to that observed under resting conditions. Only mTagBFP2/CCR3 was detected, with approximately 50% of eosinophils expressing it in the BM and 85% in the circulation (Figure 6D).

**Figure 6.**
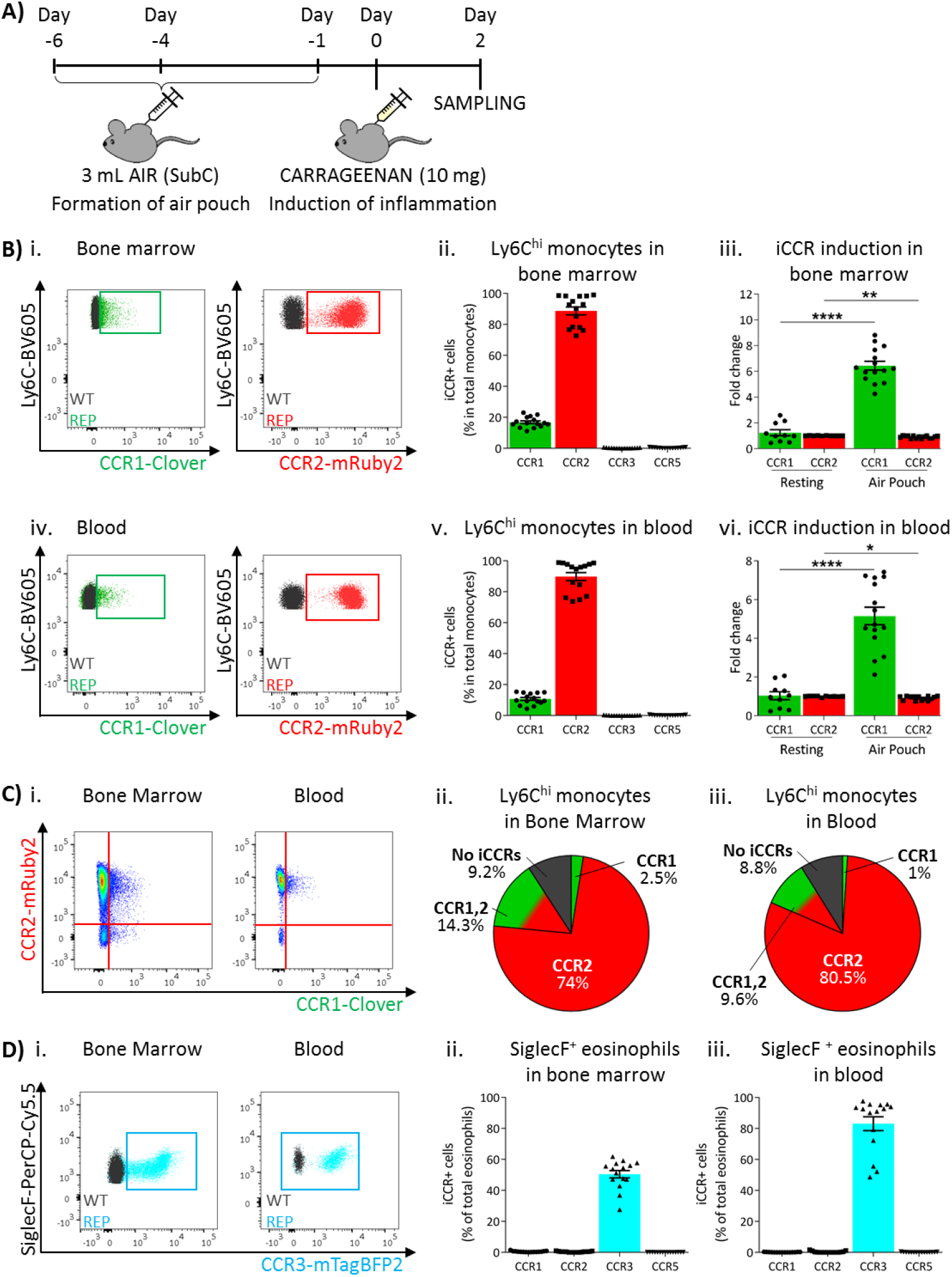
iCCR expression in acutely inflamed bone marrow and blood leukocytes. A) Schematic of the procedure used to induce acute inflammation using the air pouch model. B) Flow cytometric analysis of Clover/CCR1 and mRuby2/CCR2 expression in (i) bone marrow and (iv) circulating Ly6C^hi^ monocytes. Quantification of the percentage of (ii) bone marrow and (v) circulating Ly6C^hi^ monocytes expressing the iCCR reporters. Quantification of the fold change increase in CCR1 and CCR2 expression by (iii) bone marrow and (vi) circulating Ly6C^hi^ monocytes after induction of inflammation. C) (i) Flow cytometric analysis and distribution of Clover/CCR1 and mRuby2/CCR2 in (ii) bone marrow and (iii) circulating Ly6C^hi^ monocytes in the air pouch model. D) (i) Flow cytometric analysis of mTagBFP2/CCR3 expression in bone marrow and circulating SiglecF^+^ eosinophils. Quantification of the percentage of SiglecF^+^ eosinophils expressing the iCCR reporters in (ii) bone marrow and (iii) blood. Data on Bii, Biii, Bv, Bvi, Dii and Diii are shown as mean ±SEM (N=10 for resting mice or N=15 for carrageenan treated mice) and are compiled from at least three separate experiments. Each data point represents a measurement from a single mouse. Blots in Bi, Biv and Di are combinatorial blots showing reporter expression in iCCR REP and WT (control for background autofluorescence) mice. Normally distributed data on Biii and Bvi were analysed using unpaired t-test with or without Welch’s correction, according to their standard deviations. Not normally distributed data were analysed using Mann-Whitney or Kolmogorov-Smirnov, according to the standard deviations. *p <0.05; **p <0.01; ****p < 0.0001.

These results suggested that sustained inflammation leads to enhanced CCR1 expression on BM and circulating monocytes. We therefore used a different model to test this hypothesis. We have previously shown that CCR1 expression can be upregulated following in vitro interferon gamma (IFNγ) treatment of cells (data not shown). We therefore implanted IFNγ–loaded subcutaneous osmotic pumps (or PBS control) in iCCR-REP mice providing continuous IFNγ release into the circulation (Figure 7Ai). After 7 days, BM and blood monocytes were analysed to determine expression of Clover/CCR1. As shown in Figure 7Aii-iii, we observed a 6-fold increase in CCR1 expression in BM monocytes after IFNγ treatment. Similarly, in peripheral blood, a trend towards increased CCR1 was observed (Figure 7Aiv and v) although this was not statistically significant. In all cases, Clover/CCR1 was co-expressed with mRuby2/CCR2 in this model (Figure 7B).

**Figure 7.**
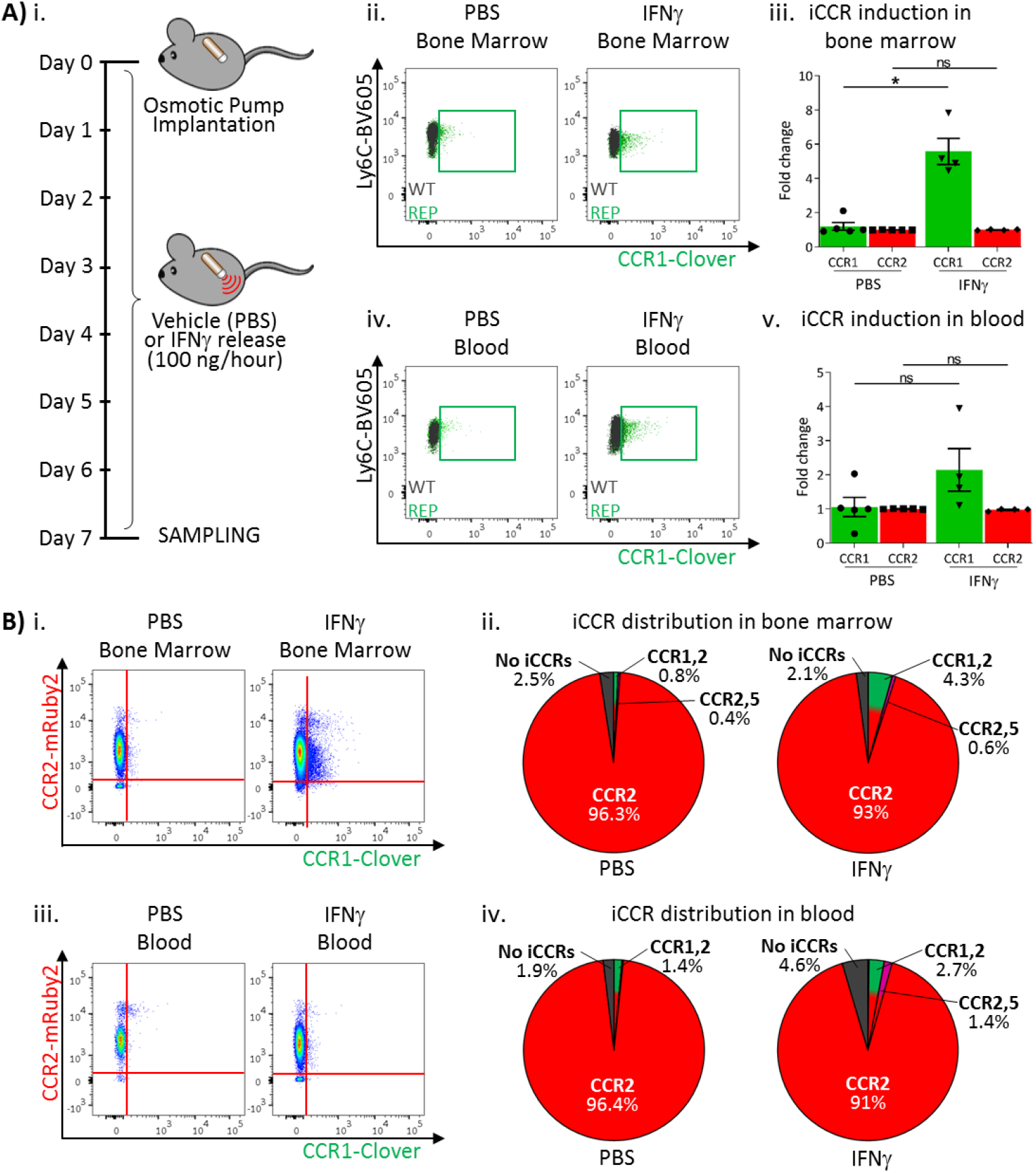
iCCR expression in bone marrow and blood under sustained inflammation. A) (i) Schematic of the procedure used to induce sustained inflammation using interferon gamma (IFNγ) release from osmotic pumps. Flow cytometric analysis of Clover/CCR1 expression in (ii) bone marrow and (iv) circulating Ly6C^hi^ monocytes. Quantification of the fold change increase in CCR1 and CCR2 expression in (iii) bone marrow and (v) circulating Ly6C^hi^ monocytes after the induction of sustained inflammation. B) Flow cytometric analysis and distribution of Clover/CCR1 and mRuby2/CCR2 in (i and ii) bone marrow and (iii and iv) circulating Ly6C^hi^ monocytes. Data on A–B are compiled from at least two separate experiments. Data on Aiii and Av are shown as mean ±SEM (N=5 for PBS-treated mice or N=4 for IFNγ-treated mice). Each data point represents a measurement from a single mouse. Blots in Aii and Aiv are combinatorial blots showing reporter expression in iCCR REP and WT (control for background autofluorescence) mice. Normally distributed data were analysed using unpaired t-test with or without Welch’s correction, according to their standard deviations. Not normally distributed data were analysed using Mann-Whitney or Kolmogorov-Smirnov, according to the standard deviations. *p <0.05; ns, not significant.

Together, these results confirm that inflammation is associated with induced expression of CCR1 in BM and circulating monocytes.

### iCCR expression in inflamed tissues: the air-pouch model

We next analysed myeloid cell iCCR expression in the inflamed air-pouch. We first examined the membrane that surrounds the inflamed cavity (gating strategies: Figure S1C). Recently infiltrated Ly6C^hi^ monocytes and CD11b^+^F480^+^ macrophages retain expression of mRuby2/CCR2 (Figure 8A), indicative of the rapid turnover of this population under inflammatory conditions. Approximately 20% of Ly6C^hi^ monocytes expressed Clover/CCR1 (Figure 8Aii), whereas expression of this receptor was much lower (Figures 4 and 5) in intra-tissue monocytes under resting conditions. Clover/CCR1 expression was further increased as Ly6C^hi^ monocytes differentiated into CD11b^+^F480^+^ macrophages, with approximately 40% of this population now expressing the receptor (Figure 8Aiii and iv). These data, together with the induction of Clover/CCR1 in inflamed BM and blood monocytes, suggest a role for this receptor in monocyte recruitment and macrophage migration in this inflammation model. In contrast, iRFP682/CCR5 expression was confined to a small fraction of the monocyte and macrophage populations (Figure 8A), whereas its expression was abundant on resting tissue macrophages (Figures 4 and 5). This suggests a less significant contribution of CCR5 to monocyte and macrophage motility in the air-pouch model.

**Figure 8.**
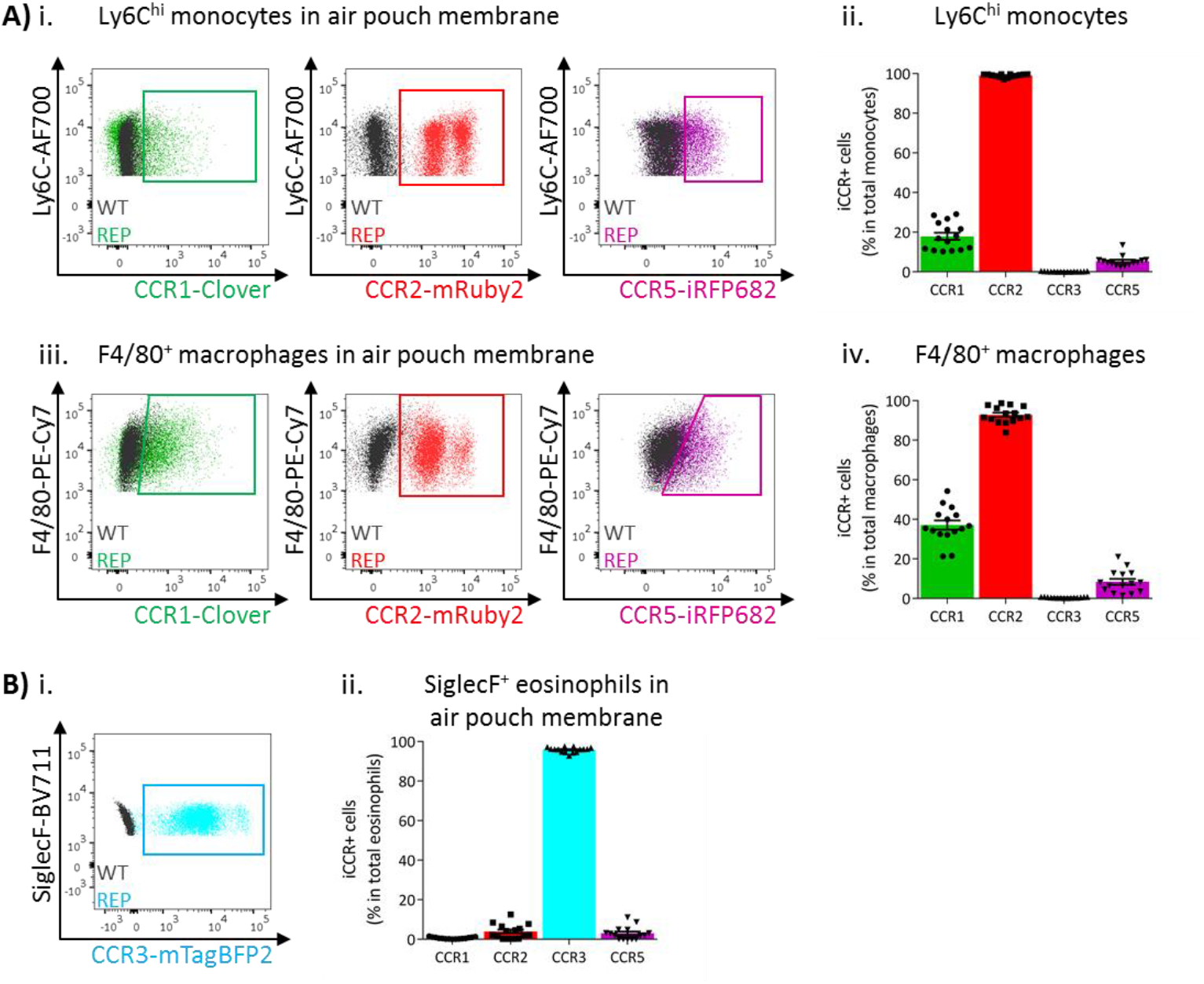
iCCR expression in inflamed tissues: the air pouch model. A) Flow cytometric analysis of Clover/CCR1, mRuby2/CCR2 and iRFP682/CCR5 expression in (i) Ly6C^hi^ monocytes and (iii) F4/80^+^ macrophages isolated from the inflamed air pouch. Quantification of iCCR reporter expression in (ii) Ly6C^hi^ monocytes and (iv) F4/80^+^ macrophages. B) (i) Flow cytometric analysis of mTagBFP2/CCR3 expression in SiglecF^+^ eosinophils isolated from the inflamed air pouch. (ii) Quantification of iCCR reporter expression in SiglecF^+^ eosinophils. Data on Aii, Aiv and Bii are shown as mean ±SEM (N=15) and are compiled from at least three separate experiments. Each data point represents a measurement from a single mouse. Blots in Ai, Aiii and Bi are combinatorial blots showing reporter expression in iCCR REP and WT (control for background autofluorescence) mice.

As shown in Figure 8B, eosinophils in the air-pouch retain exclusive expression of mTagBFP2/CCR3, confirming the importance of this receptor for eosinophil recruitment to inflamed sites. We did not detect expression of any iCCR reporter in neutrophils (data not shown).

### iCCR expression in inflamed tissues: the intranasal LPS model

To determine if the pattern of iCCR expression detected in the air-pouch model was also seen in a more relevant inflammatory model, we next used an LPS model of lung inflammation. Here, LPS (or vehicle) was administered intranasally to iCCR-REP mice or WT littermates (Figure 9Ai). 48 hours later, inflamed lungs were dissected and myeloid cell content examined (gating strategies: Figure S2A). Infiltration of Ly6C^hi^ monocytes and CD11b^+^F480^+^ macrophages into inflamed lung was confirmed by flow cytometry (Figure 9Aii). Consistent with the findings from the air-pouch model, Ly6C^hi^ monocytes and CD11b^+^F480^+^ macrophages both retain expression of mRuby2/CCR2, indicative of their recent infiltration into the inflamed lung (Figure 9Aiii-iv). Similarly, 23% of the lung monocyte population expressed Clover/CCR1 after LPS treatment, compared to only 3.5% of monocytes in vehicle treated mice (Figure 9Aiii). This level of expression is maintained in mature CD11b^+^F480^+^ macrophages from LPS-treated lungs (Figure 9Aiv, B and C), confirming the rapid induction of CCR1 in infiltrated Ly6C^hi^ monocytes. Finally, also consistent with observations in the air-pouch model, iRFP682/CCR5 was detected in only 7.5% of Ly6C^hi^ monocytes and 12% of CD11b^+^F480^+^ macrophages after LPS administration, whereas 65% of CD11b^+^F480^+^ macrophages expressed it in the vehicle-treated lungs (Figure 9Aiii-iv, B and C). These data suggest that resident CCR5^+^ macrophages are rapidly replaced with CCR1^+^ macrophages after induction of inflammation and indicate a less significant contribution of CCR5 to monocyte and macrophage recruitment and migration in the early stages of the inflammatory response to LPS.

**Figure 9.**
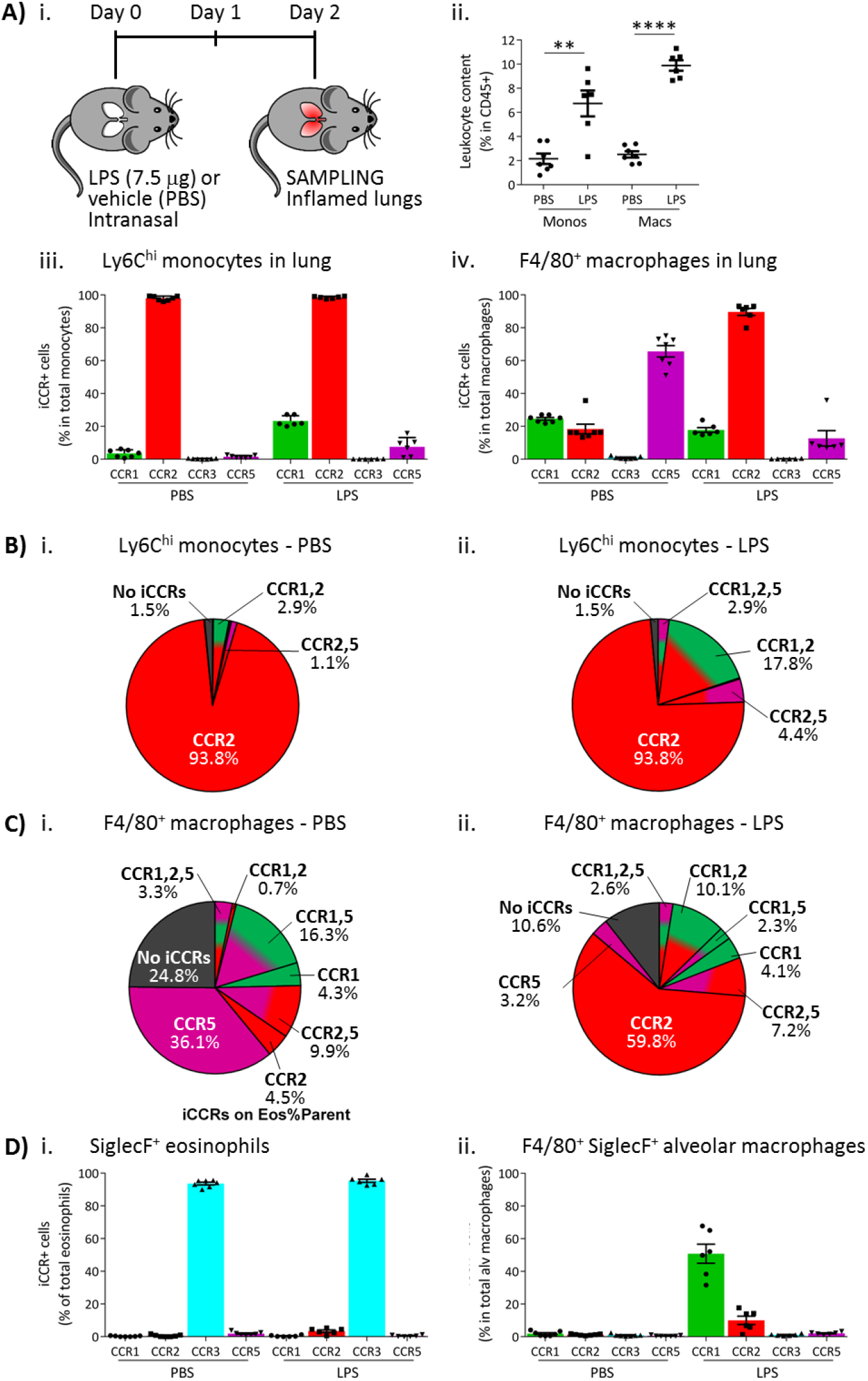
iCCR expression in inflamed tissues: the intranasal LPS model. A) (i) Schematic of the procedure used to induce acute lung inflammation using intranasal administration of LPS. (ii) Quantification of monocyte and macrophage infiltration into the inflamed lungs. Quantification of iCCR reporter expression in (iii) Ly6C^hi^ monocytes and (iv) F4/80^+^ macrophages isolated from lungs of vehicle (PBS) and LPS-treated mice. B) Distribution of Clover/CCR1, mRuby2/CCR2 and iRFP682/CCR5 on Ly6C^hi^ monocytes isolated from lungs of (i) vehicle (PBS) and (ii) LPS-treated mice. C) Distribution of Clover/CCR1, mRuby2/CCR2 and iRFP682/CCR5 on F4/80^+^ macrophages isolated from lungs of (i) vehicle (PBS) and (ii) LPS-treated mice. D) Quantification of iCCR reporter expression on (i) SiglecF^+^ eosinophils and (ii) SiglecF^+^ F4/80^+^ alveolar macrophages isolated from lungs of vehicle and LPS-treated mice. Data in A–D are compiled from at least two separate experiments. Data on Aii, Aiii, Aiv, Di and Dii are shown as mean ±SEM (N=7 for vehicle-treated mice or N=6 for LPS-treated mice). Each data point represents a measurement from a single mouse. Data on Aii were analysed using unpaired t-test. **p <0.01; ****p < 0.0001. Abbreviations are: Monos, monocytes; Macs, macrophages.

We also evaluated iCCR expression on SiglecF^+^ eosinophils and SiglecF^+^F480^+^ alveolar macrophages after LPS inoculation. As expected, eosinophils showed exclusive expression of mTagBFP2/CCR3 (Figure 9Di), confirming the importance of this receptor for their recruitment into the inflamed site. Alveolar macrophages do not express any iCCR after vehicle treatment (resting conditions). However, after LPS administration, 50% of the population shows expression of Clover/CCR1 (Figure 9Dii), suggesting that this receptor is also important for their intra-tissue function under inflammatory conditions. We did not detect expression of iCCR reporters in neutrophils from inflamed lungs (data not shown).

## Discussion

The iCCRs are responsible for the mobilisation, recruitment and intra-tissue dynamics of all non-neutrophilic myeloid cell subsets as well as some lymphoid subsets. Analysis of their in vivo expression dynamics has been hampered by the difficulties associated with generating combinatorial reporter mice using individual reporter strains(Hirai et al., 2014; Luckow et al., 2009; Saederup et al., 2010), due to their genomic proximity and the incompatibility of the reporters used in these strains. We now report the generation, and analysis, of transgenic mice expressing spectrally-distinct fluorescent reporters for each of the four iCCRs. To our knowledge, this is the first mouse model allowing simultaneous, and combinatorial, analysis of four different fluorescent reporters for the study of specific protein expression, and provides a template for the generation of similar reporter mice covering other functionally-linked genomic loci.

The iCCR-REP mice are viable, display normal leukocyte trafficking and, importantly, reporter expression accurately reflects surface iCCR presentation. Antibodies are widely used to study iCCR expression. However, our results demonstrate that these antibodies show background non-specific staining due to the high degree of homology between different iCCRs. Also in our hands, and as described by others(Hirai et al., 2014), commercially available antibodies for CCR1 do not bind efficiently to the receptor. In contrast, our iCCR-REP mice display highly specific expression of the reporters, representing a powerful tool for the flow cytometric, and imaging-based, analysis of in vivo iCCR expression.

Importantly, the analysis of the iCCR-REP mice has allowed us, for the first time, to unequivocally establish iCCR expression patterns on myeloid cells at rest and during inflammatory responses. In contrast to previous reports of multiple receptor expression by individual leukocyte subtypes(Haringman, Smeets, Reinders-Blankert, & Tak, 2006; Tacke et al., 2007; Weber et al., 2000), our data indicate that iCCR expression on individual cell subsets in resting mice is selective. Thus, most Ly6C^hi^ monocytes in BM and blood exclusively express CCR2, with only a minor fraction of these cells co-expressing CCR1 and CCR2. Eosinophils only express CCR3. Tissue-resident macrophages downregulate CCR2 and express CCR1 and high levels of CCR5 either alone or in combination. This suggests a hierarchical relationship in which monocytes use CCR2 to egress from BM and infiltrate resting tissues but upregulate CCR1 and CCR5 upon differentiation to macrophages or moDCs. This model was further confirmed by our in vitro studies using GM-CSF BM derived moDCs. While Ly6C^hi^ monocytes freshly isolated from BM expressed almost exclusively CCR2, culture in GM-CSF containing media induced gradual upregulation of CCR1 and CCR5. After 9 days in culture, monocytes were fully differentiated into moDCs and expressed high levels of CCR5. We propose that CCR1 and CCR5 are predominantly involved in intra-tissue migration and not recruitment from the circulation.

Under inflammatory conditions, BM and circulating Ly6C^hi^ monocytes still express CCR2. However, the fraction co-expressing CCR1 increases. We reasoned that this may be a consequence of elevated systemic inflammatory cytokines and indeed sustained increase in systemic IFNγ levels recapitulated this phenotype. As the iCCRs, and other chemokine receptors, are transcriptionally regulated in response to a variety of cytokines, this raises the possibility that reports of multiple chemokine receptor expression by inflammatory cells in patients with immune and inflammatory disorders, is not a reflection of homeostasis but a response to the systemically inflamed environment in these patients. We speculate that this may provide alternative options for leukocyte recruitment from the periphery to inflamed sites and contribute to the failure of specific receptor-targeting therapeutics in inflammatory disease. This hypothesis is supported by our previous findings showing that, in the absence of CCR2, a small proportion of Ly6C^hi^ monocytes can still infiltrate into inflamed tissues(Dyer et al., 2019).

After infiltration into the inflamed site, Ly6C^hi^ monocytes still express CCR2 and eosinophils CCR3. Whether these populations represent functionally distinct subsets remains unclear. Recently infiltrated Ly6C^hi^ monocytes also rapidly increase CCR1 expression, which is maintained after differentiation into F480^+^ macrophages. This suggests that, in an inflamed context, CCR1 has a pivotal role in directing intra-tissue migration of monocytes and macrophages towards the focus of inflammation and supports the hypothesis that circulating monocytes co-expressing CCR1 and CCR2 might have a recruitment advantage over CCR2 only monocytes. This is further supported by the fact that alveolar macrophages in lung also upregulate CCR1 expression under inflamed conditions, whereas this population does not express any of the iCCRs under resting conditions.

Interestingly, our analyses show that CCR5 is preferentially expressed by macrophages in resting tissues, whereas CCR1 is strongly associated with macrophages recently recruited to inflamed tissues. This could be explained by a hierarchical model of iCCR expression, where bone marrow and circulating Ly6C^hi^ monocytes express CCR2 but, immediately after extravasation, they downregulate CCR2 and upregulate CCR1 as they differentiate into F480^+^ macrophages. CCR1 would direct macrophages in the early stages of intra-tissue migration. Finally, F480^+^ macrophages would induce expression of CCR5 and slowly downregulate CCR1 as they fully differentiate into mature macrophages. This model would explain the absence of macrophages expressing exclusively CCR1 in resting tissues, whereas a large proportion of them are CCR1^+^CCR5^+^ or CCR5^+^ only. Alternatively, CCR1 and CCR5 might be expressed in response to different inflammatory stimuli or have different functions in the inflamed tissue. We used carrageenan and LPS to induce inflammation in our study, both signalling through toll-like receptor 4 (TLR-4)(Myers, Deaver, & Lewandowski, 2019; Solov’eva, Davydova, Krasikova, & Yermak, 2013). This could explain their similar responses, mediated by CCR1^+^ macrophages. However, CCR5 has been reported to be essential for macrophage recruitment to virus-infected tissues(Glass et al., 2005), raising the possibility that alternative inflammatory stimuli could trigger differential responses.

Finally, we failed to detect expression of any of the iCCRs in neutrophils either at rest or during inflammation. We analysed three different founder lines for the iCCR REP mice and obtained identical results with all of them. This contrasts with previous reports(Fujimura et al., 2015; Gao et al., 1997; Gerard et al., 1997) indicating roles for CCR1 in neutrophil recruitment, but is supported by our findings with iCCR deficient mice, that show that absence of these chemokine receptors does not lead to deficiencies in neutrophil recruitment to resting or inflamed tissues(Dyer et al., 2019). It is possible that differences in inflammatory models used in the various studies, or animal housing arrangements, may have contributed to this apparent disagreement with the literature.

In summary, therefore, our analyses indicate that leukocyte iCCR expression is more specific (summarised in Figure S6) than previously believed, suggesting non-redundant roles for the iCCRs in leukocyte cell trafficking at rest and in acute inflammation. We propose that the iCCR-REP mice represent a transformational technical advance permitting in-depth analysis of receptor expression in a range of resting and pathological conditions that will shed important light on chemokine receptor biology and potentially inform future rational drug design.

## Acknowledgements

We acknowledge the assistance of the Institute of Infection, Immunity and Inflammation Flow Core Facility at the University of Glasgow. We also thank Dr. Michael Z. Lin and Prof. Vladislav V. Verkhusha for their advice on the selection of the fluorescent reporters. Work in GJG’s laboratory is funded by a Wellcome Trust Investigator Award (REF: 217093/Z/19/Z) and an MRC Programme Grant (MRV0109721).

## Author contributions

LM-R, RB, DD, FV, CH, FS, SL and JF performed experiments. GJG and AFS conceived the study. All authors were involved in the writing of the manuscript and the analysis of the data. All authors approved the final manuscript.

## Data and materials availability

This study includes no data deposited in external repositories. All datasets generated or analysed during the current study are included in this published article (and its supplementary information files). Raw data will be made available on reasonable request by the corresponding authors, Gerard J Graham or Laura Medina-Ruiz. Mouse lines generated in this study will be available to non-commercial organisations on request.

## Materials and Methods

### Mouse generation and maintenance

The iCCR-REP mice were generated in collaboration with Taconic Biosciences. First, the *iCcrs* were targeted in a BAC using counterselection recombineering as previously described(Wang et al., 2014) in order to replace the coding sequence of each one of the *iCcrs* with the sequence coding for a different fluorescent reporter. Then, the iCCR-REP cluster was incorporated into the mouse genome using pro-nuclear injection, thus generating transgenic iCCR-REP mice. The presence of the iCCR-REP cluster in these animals was confirmed by PCR using primers specific for the iCCR reporters (Table S1A). Quantification of the number of copies of the iCCR-REP cluster inserted into the mouse genome was done by QPCR with the primers listed in Table S1B, using the TBP gene as a reference, because it sits outside of the iCCR cluster and remained unaltered in the process. Finally, the iCCR-REP cluster was localised to chromosome 16:82389380-82392016 using targeted locus amplification (TLA) as described elsewhere(de Vree et al., 2014).

iCCR-def mice were generated in-house(Dyer et al., 2019) and CCR5-def mice were a kind gift from Dr. Takanori Kitamura, University of Edinburgh. All animal strains were generated and maintained on a C57BL/6 background and were bred in a specific pathogen free environment in the animal facility of the Beatson Institute for Cancer Research (Glasgow). Routine genotyping of all animals was done by PCR of ear samples (Transnetyx). All experiments were done on animals 10-12 weeks old, using congenic WT animals derived from heterozygous crosses as controls. All experiments were carried out under the auspices of a UK Home Office Project License and were approved by the local Ethical Review Committee.

### Resting tissue isolation

Resting mice were sacrificed and tissues extracted for flow cytometry or RNA analysis. Blood was extracted from the vena cava and mixed with 100 μL of 0.5 M EDTA (Thermo Fisher Scientific) prior to red blood cell lysis using an ACK lysis solution (Thermo Fisher Scientific) as per manufacturer’s instructions. Leukocyte content was then used for further analysis. Immediately after blood isolation, mice were perfused using 20 mL of PBS (Thermo Fisher Scientific) containing 2 mM EDTA (Thermo Fisher Scientific).

Bone marrow was then extracted from the femur and tibia of the mice, red blood cells lysed as described above and leukocyte content used in downstream analyses.

Spleen, lungs and kidneys were isolated from perfused animals and incubated over-night (O/N) in RNAlater™ stabilisation solution (Thermo Fisher Scientific) for RNA analysis. Alternatively, these tissues were processed for antibody staining and flow cytometry analysis. Dissected spleens were filtered through 70 μm nylon mesh membranes, washed with PBS and red blood cells lysed as described above, prior to antibody staining. Lungs were cut into small pieces and digested in 5 mL of RPMI containing DNase I (100 μg/mL, Roche), Dispase II (800 μg /mL, Roche) and Collagenase P (200 μg/mL, Roche) for 90 minutes at 37°C. Kidneys were cut into small pieces and digested in 4 mL of PBS containing calcium and magnesium, HEPES (20 mM, VWR), Collagenase I (1.8 mg/mL, Sigma-Aldrich), Collagenase XI (156 μg/mL, Sigma-Aldrich) and Hyaluronidase (158 μg/mL, Sigma-Aldrich) for 20 minutes at 37°C. After digestion, enzymes were deactivated using 20 μL of foetal bovine serum (FBS) and lung and kidney cell suspensions were filtered through 70 μm nylon mesh membranes, washed with PBS and stained for flow cytometry analysis.

### Air pouch model

The air pouch model of inflammation was used as previously described(Dyer et al., 2019). In brief, 3 mL of sterile air were injected under the dorsal skin on 3 occasions over a period of 6 days to induce the formation of a subcutaneous air cavity. 1 mL of sterile carrageenan (1% (w/v) in PBS, Sigma-Aldrich) was inoculated into the cavity to induce inflammation 24 h after the last air injection. 48 hours later, mice were culled, and blood and bone marrow samples were collected as detailed above. The cavity was flushed with 3 mL of PBS containing 2 mM EDTA and 2% (v/v) FBS and the lavage fluid was collected, washed with PBS and stained for flow cytometry analysis. The membrane surrounding the air pouch was then isolated and digested in 1 mL of Hanks buffered saline solution (HBSS, Thermo Fisher Scientific) containing 0.44 Wünsch units of Liberase (Roche) for 1 hour at 37°C and 1000 rpm shaking. After digestion, Liberase was neutralised using 20 μL of FBS and cell suspensions were filtered through 70 μm nylon mesh membranes, washed with PBS and stained for flow cytometry analysis.

### LPS model of lung inflammation

For the induction of lung inflammation using *E. coli* lipopolysaccharide (LPS), mice were anaesthetised using inhaled isoflurane (4% (v/v) isoflurane and 2 l O_2_/min) and 30 μl of LPS (250 μg/mL, Sigma-Aldrich) or vehicle PBS were administered intranasally. 48 hours later, mice were culled, perfused and lungs were isolated for flow cytometry analysis as detailed above.

### Implantation of IFNγ–loaded subcutaneous osmotic pumps

Mice were anaesthetised using inhaled isoflurane (2% (v/v) isoflurane and 2 l O_2_/min) and maintained under these conditions during the surgical procedure. A small pocket was generated under the dorsal skin, where IFNγ- (100 ng/μL) or vehicle PBS-loaded osmotic pumps (Alzet® osmotic pumps, model 2001; Charles River) were implanted. Infusion of IFNγ (100ng/hour) or PBS was maintained for 7 days. After this time, animals were sacrificed and bone marrow and blood were extracted for flow cytometry analysis as described above.

### Flow cytometry

Tissue lysates were prepared as described above and stained for 20 minutes at 4°C with 100 μl of fixable viability stain (eBioscience). Cells were then washed in FACS buffer (PBS containing 2 mM EDTA and 2% (v/v) FBS) and stained for 20 minutes at 4°C with 50 μl of antibody cocktail containing subset-specific antibodies (Supplementary Table 2) diluted in Brilliant Stain Buffer (BD Biosciences). Cells were washed in FACS buffer and fixed for 20 minutes at 4°C in 100 μl of Fixation Buffer (BioLegend).

Stained samples were analysed on a BD LSRFortessa™ flow cytometer (BD Biosciences) and data analysis was performed using FlowJo software (FlowJo).

### Generation of bone marrow derived moDCs

Bone Marrow cells were isolated as described above and 10^7^ cells were resuspended in 10 mL of RPMI-1640 (Sigma) supplemented with FBS (10% v/v), L-glutamine (1% v/v), 50μM β-mercaptoethanol, primocin and 20 ng/mL of murine recombinant GM-CSF (Peprotech). Cells were transferred to tissue culture treated petri dishes. At Days 2 and 9, the medium was collected, centrifuged at 300 *g* for 5 minutes and pelleted cells were analysed for the expression of iCCR reporters by flow cytometry. Monocytes and moDCs were identified on the basis of Ly6C, F480, CD11c, and MHCII expression (monocytes were CD11c^-^ F480^+^ Ly6C^hi^; moDC precursors were F480^+^ CD11c^+^ Ly6C^-^; mature moDCs were CD11c^+^ MHCII^hi^).

### RNAscope for the detection of CCR1 mRNA

Kidneys were isolated from resting mice after PBS perfusion and fixed O/N in 20 mL of 10% Neutral Buffered Formalin (Leica). Fixed kidneys were paraffin-embedded using a Shandon Citadel 1000 tissue processor (Thermo Fisher Scientific) and stored at room temperature (RT) until used. The day prior to CCR1 mRNA analysis, kidneys were wax-embedded and sliced into 6 μm sections onto SuperFrost Plus™ adhesion slides (Thermo Fisher Scientific). Slides were air-dried O/N at RT and, the following day, CCR1 mRNA was detected using an RNAscope® target probe specific for the gene with the RNAscope® 2.5 HD Reagent Kit-RED (Advanced Cell Diagnostics). Manufacturer’s instructions were followed, with minor modifications. Specifically, kidney sections were incubated in the RNAscope® Target Retrieval Reagent for 18 minutes and were treated with the RNAscope® Preotease Plus Reagent for 35 min. Images were acquired on an Evos FL Auto 2 microscope (Thermo Fisher Scientific).

### Imaging

Lungs, spleens and kidneys were isolated from resting mice after PBS perfusion. They were immersed in 4 mL of 4% paraformaldehyde (VWR), incubated at RT for 1 hour and finally incubated ON at 4°C to achieve full fixation. Tissues were then immersed in increasing concentrations of sucrose (10%, 20% and 30% (w/v) in PBS; Fisher Chemicals) for 18 to 24 hours in each solution. Finally, tissues were frozen in O.C.T.™ embedding media (Tissue-Tek) and stored at −80°C.

48 to 24 hours prior to sectioning, tissues were transferred to −20°C. After −20°C incubation, tissues were sliced into 8 μm sections onto SuperFrost Plus™ adhesion slides and transferred to −20°C again until processed for imaging.

The day of imaging, tissue sections were incubated at 60°C for 10 minutes and washed twice in PBS for 5 minutes. Slides were then incubated for 20 minutes in 0.1 M glycine (VWR) with mild shaking to reduce background autofluorescence. After glycine incubation, sections were washed 5 times with PBS for 5 minutes and then immersed in cold water until mounted using 10 μl of Mowiol mounting medium. Tissues were imaged using an Axio Imager M2 microscope (Zeiss) or a spinning disk Axio Observer Z1 confocal microscope (Zeiss).

### Quantification and statistical analysis

All analyses were performed using the Prism software package (GraphPad). Normally distributed data were analysed using unpaired t-test with or without Welch’s correction, according to their standard deviations. Not normally distributed data were analysed using Mann-Whitney or Kolmogorov-Smirnov, according to their standard deviations. In all analysis, p=0.05 was considered the limit for statistical significance.

## Supplementary Figures and Tables

**Figure S1.**
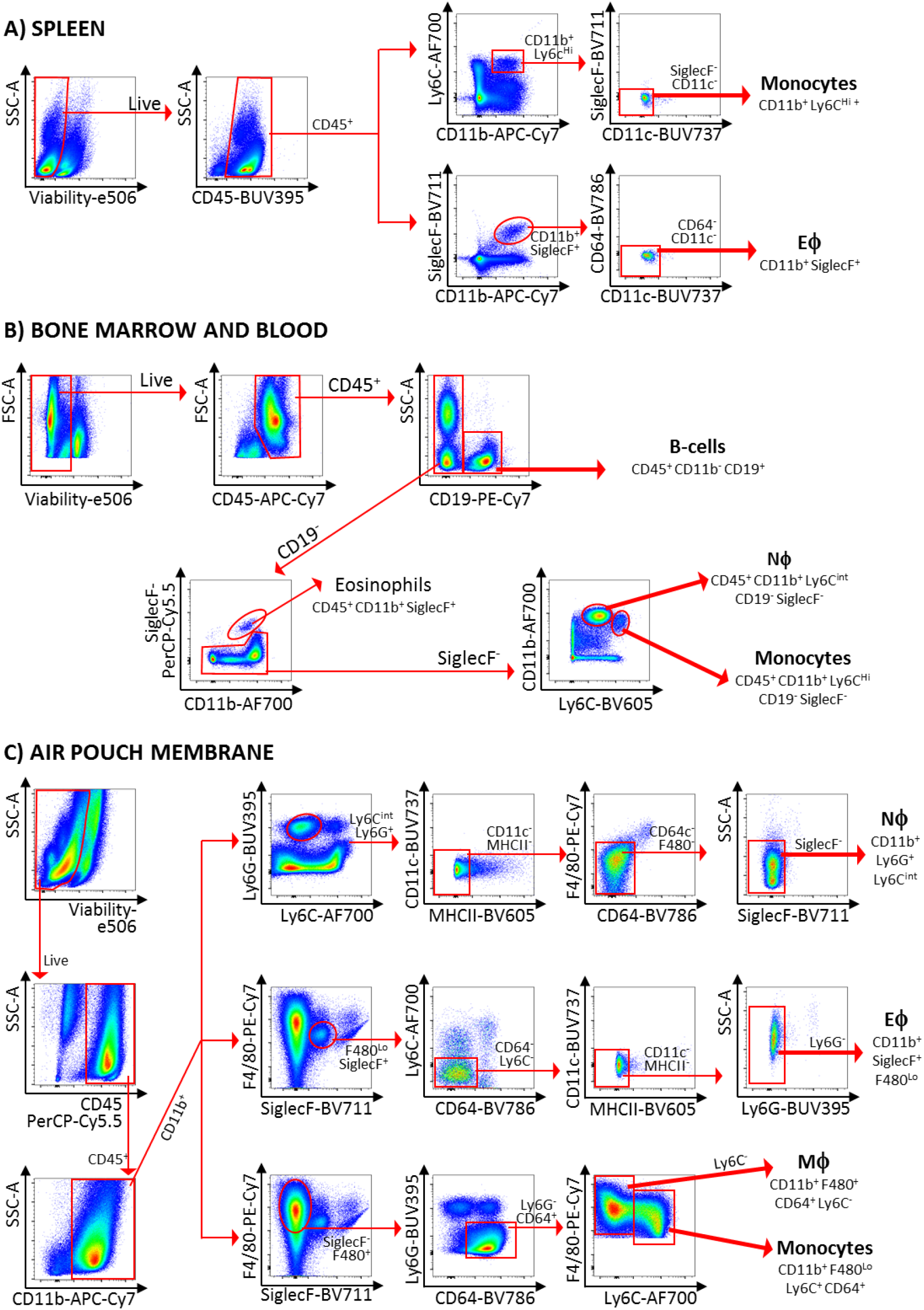
Gating strategies used in the study. Gating strategies used for the isolation of cell subsets in (A) spleen, (B) bone marrow and blood and (C) air pouch membrane. Abbreviations are: Eϕ, eosinophils; Nϕ, neutrophils; Mϕ, macrophages.

**Figure S2.**
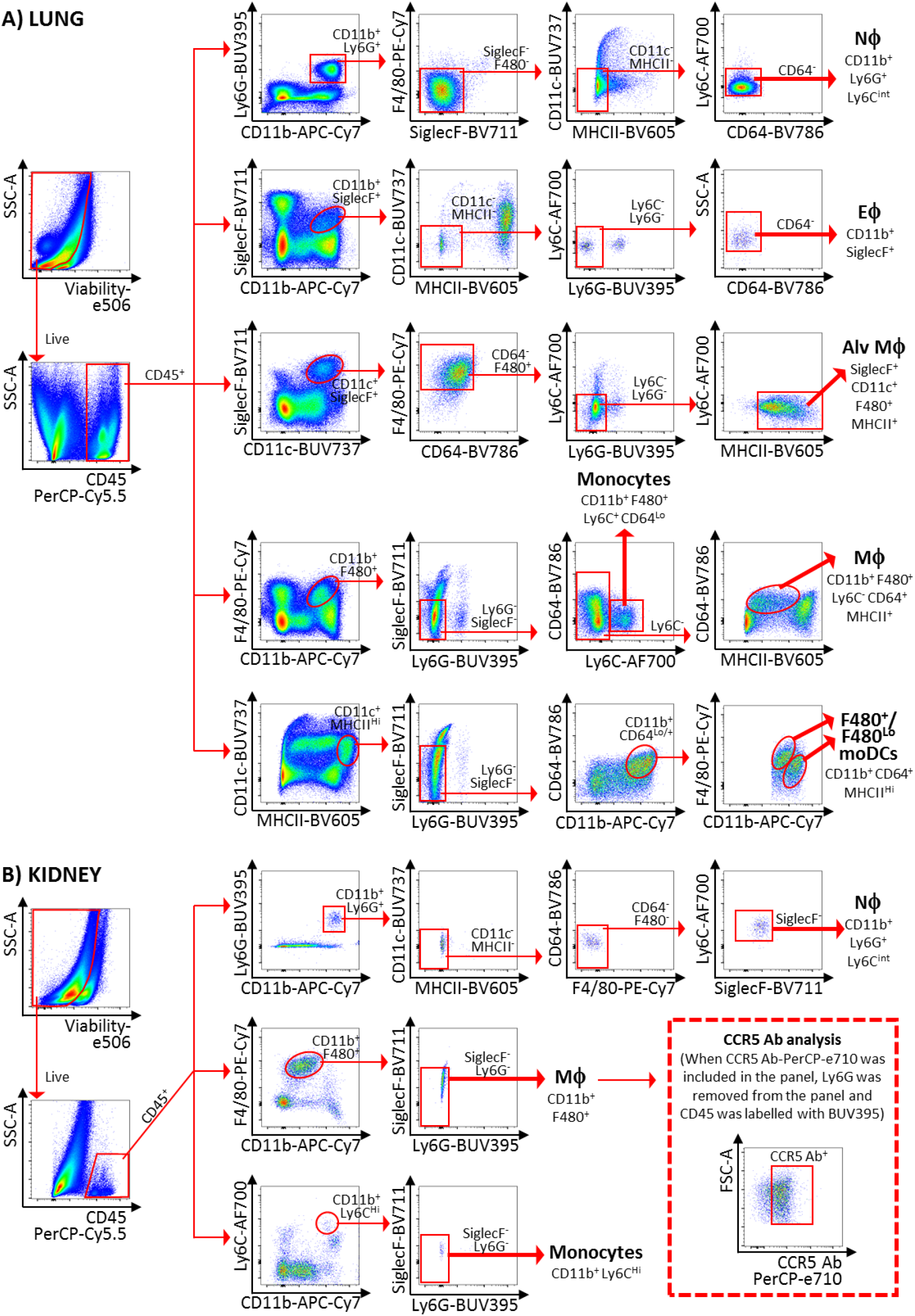
Gating strategies used in the study. Gating strategies used for the isolation of (A) lung and (B) kidney cell subsets. Abbreviations are: Nϕ, neutrophils; Eϕ, eosinophils; Alv Mϕ, alveolar macrophages; Mϕ, macrophages; moDCs, monocyte-derived dendritic cells

**Figure S3.**
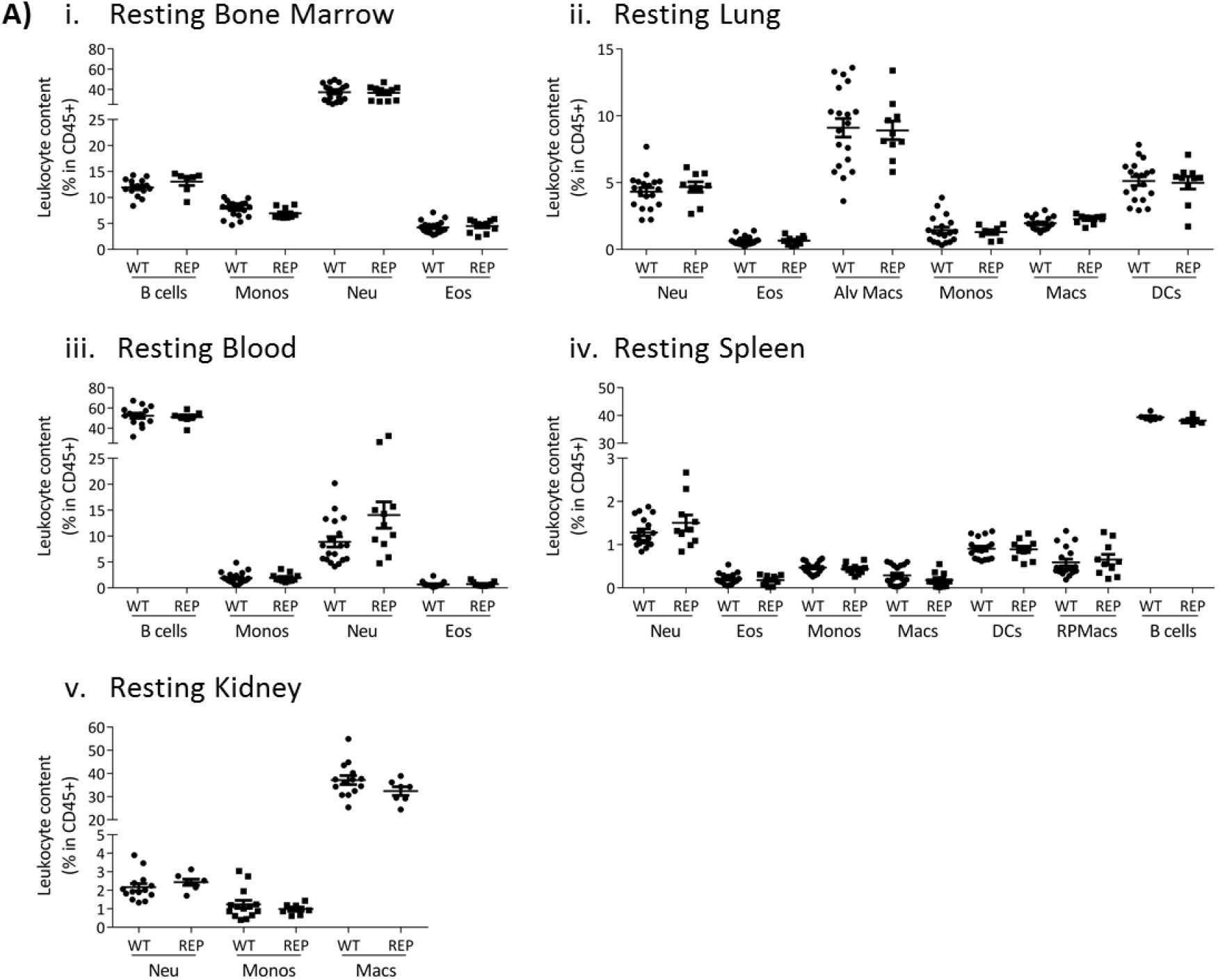
Normal leukocyte recruitment in the reporter mice. A) Leukocyte counts determined by flow cytometry in resting (i) bone marrow, (ii) lung, (iii) blood, (iv) spleen and (v) kidney. Data are shown as mean ±SEM and are compiled from at least two separate experiments. Normally distributed data were analysed using unpaired t-test with or without Welch’s correction, according to their standard deviations. Not normally distributed data were analysed using Mann-Whitney or Kolmogorov-Smirnov, according to their standard deviations. Each data point represents a measurement from a single mouse. Abbreviations are: Monos, monocytes; Macs, macrophages; Neu, neutrophils; Eos, eosinophils; DCs, dendritic cells; RPMacs, red pulp macrophages; Alv macs, alveolar macrophages.

**Figure S4.**
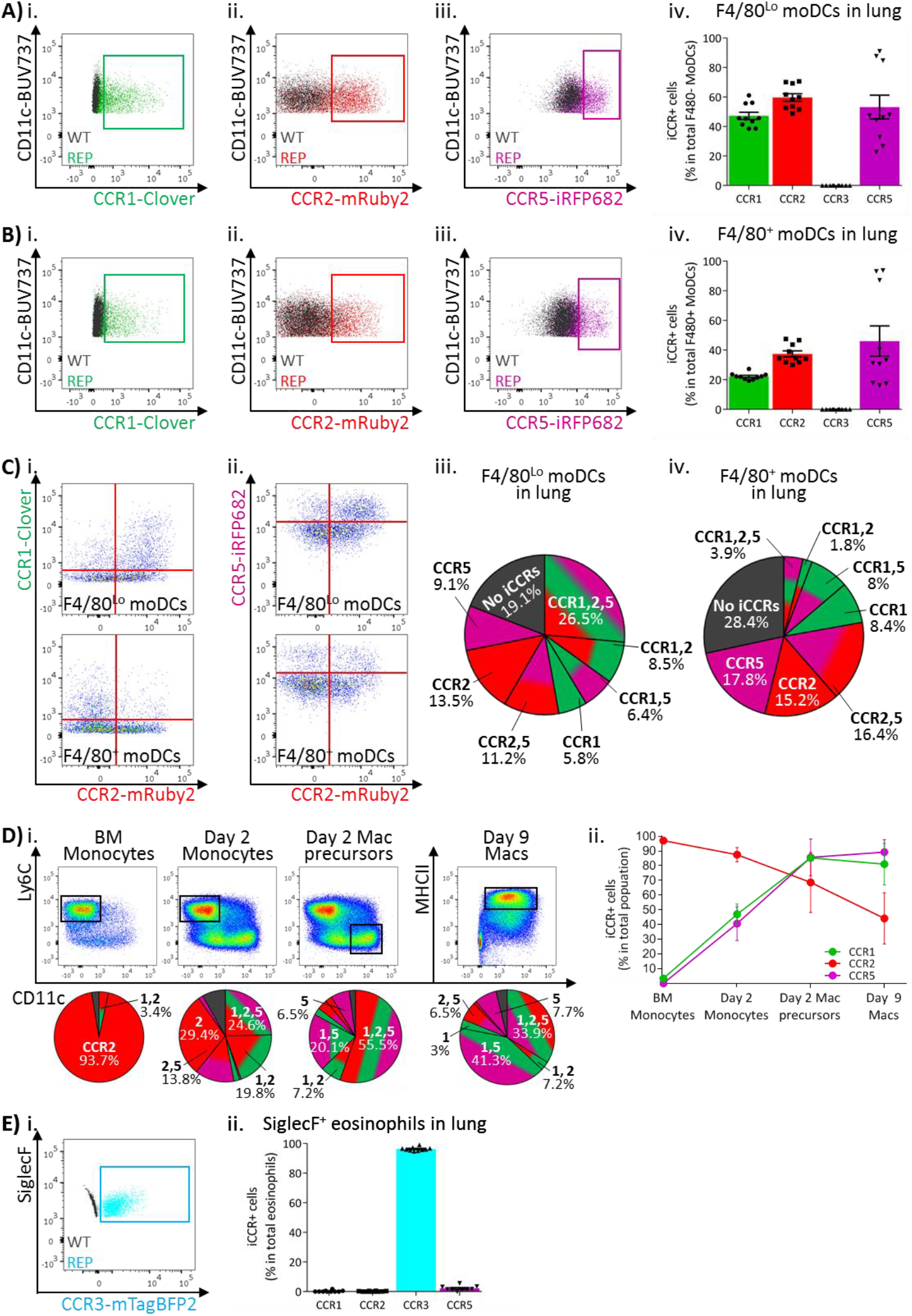
iCCR expression in resting lung. A) Flow cytometric analysis of (i) Clover/CCR1, (ii) mRuby2/CCR2 and (iii) iRFP682/CCR5 expression in F4/80^Lo^ monocyte-derived dendritic cells (moDCs) in resting lung. (iv) Quantification of the percentage of F4/80^Lo^ moDCs expressing the iCCR reporters. B) Flow cytometric analysis of (i) Clover/CCR1, (ii) mRuby2/CCR2 and (iii) iRFP682/CCR5 expression on F4/80^+^ moDCs in resting lung. (iv) Quantification of the percentage of F4/80^+^ moDCs expressing the iCCR reporters. C) Flow cytometric analysis of the co-expression of (i) Clover/CCR1 with mRuby2/CCR2 or (ii) iRFP682/CCR5 with mRuby2/CCR2 on moDCs in resting lung. Distribution of the iCCR reporters in (iii) F4/80^Lo^ and (iv) F4/80^+^ moDCs. D) Flow cytometric analysis (i) and quantification (ii) of iCCR expression on BM derived GM-CSF moDCs. E) (i) Flow cytometric analysis of mTagBFP2/CCR3 expression in SiglecF^+^ eosinophils in resting lung. (ii) Quantification of the percentage of SiglecF^+^ eosinophils expressing the iCCR reporters. Data in A–C and E are compiled from at least three separate experiments. Data in D is pooled from 5 mice. Data in Aiv, Biv and Eii are shown as mean ±SEM (N=10). Each data point represents a measurement from a single mouse. Blots in Ai, Aii, Aiii, Bi, Bii, Biii and Ei are combinatorial blots showing reporter expression in iCCR REP and WT (control for background autofluorescence) mice.

**Figure S5.**
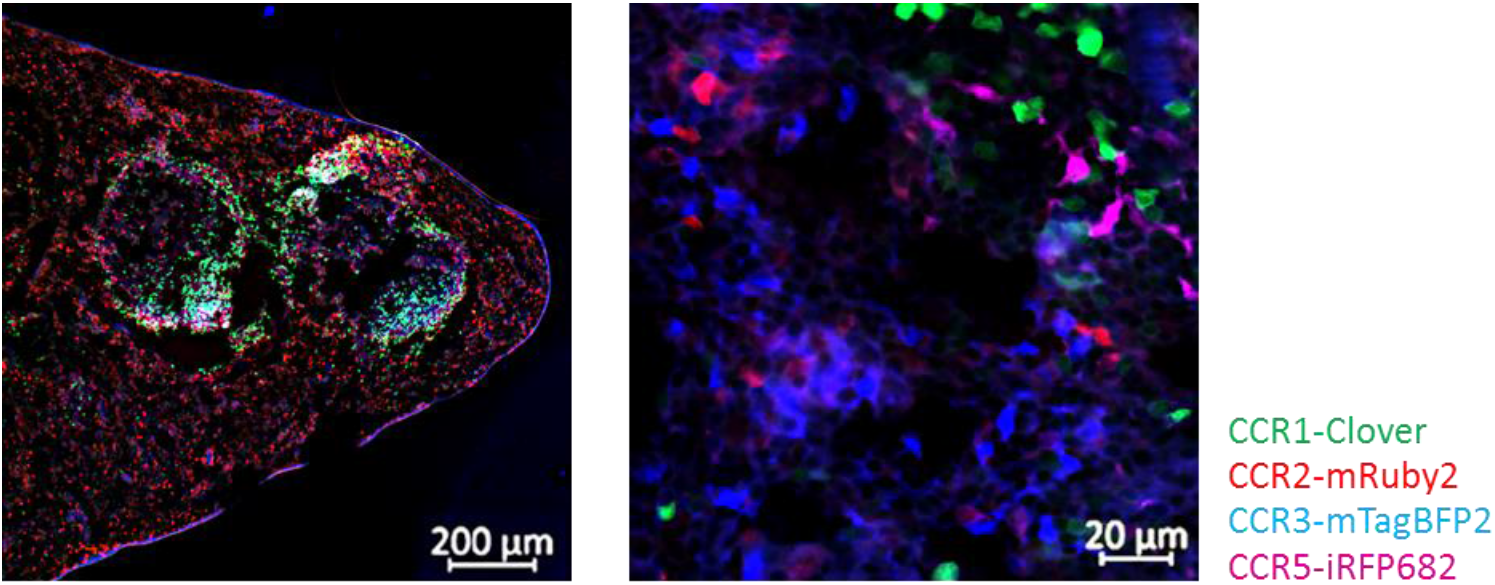
iCCR reporters are readily visualised in tissues. Spleens from resting mice were isolated and imaged using an AxioImager M2 microscope (Zeiss). Different magnifications are shown.

**Figure S6.**
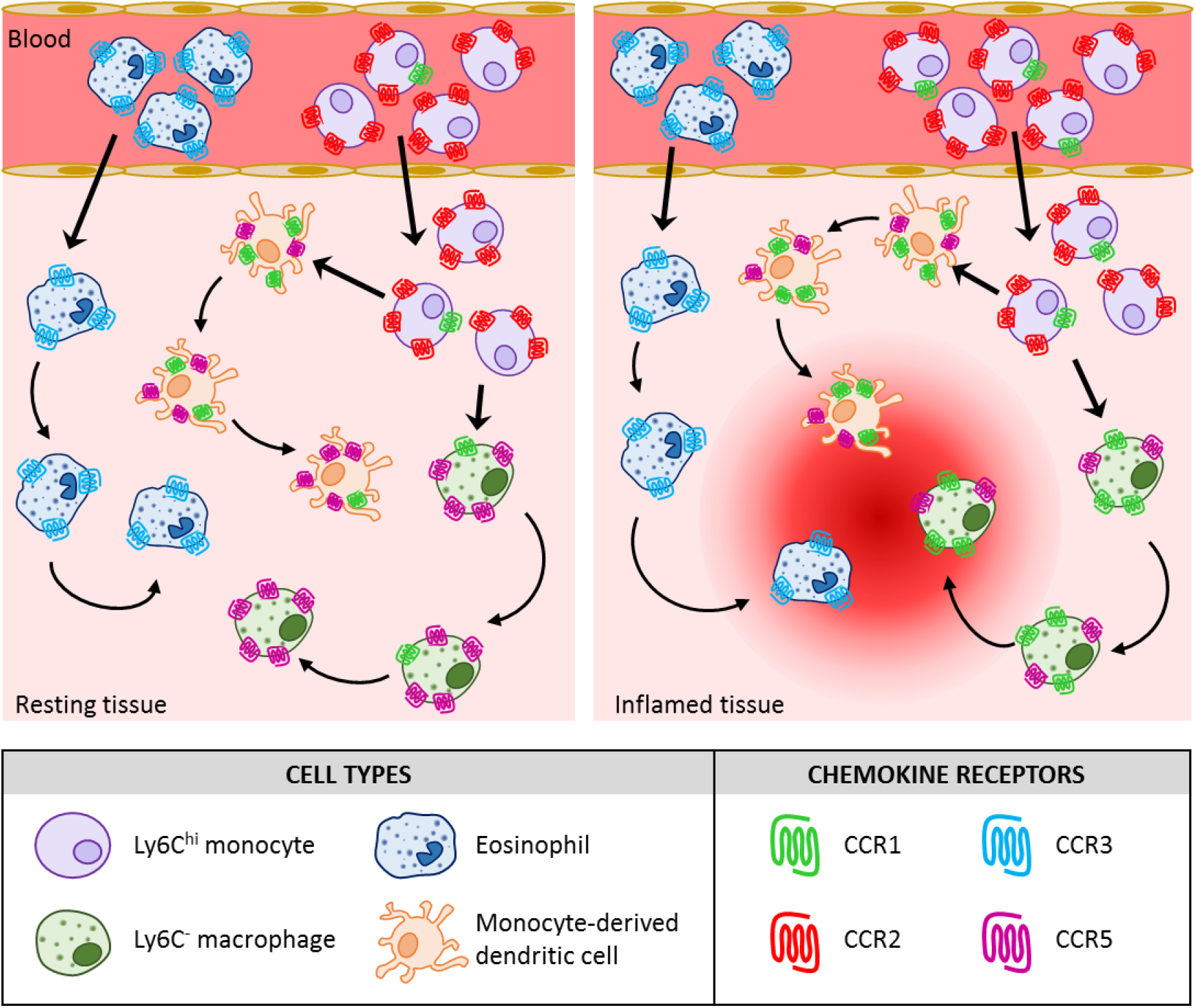
Proposed model for the contribution of the iCCRs to leukocyte migration.

**Table S1.**
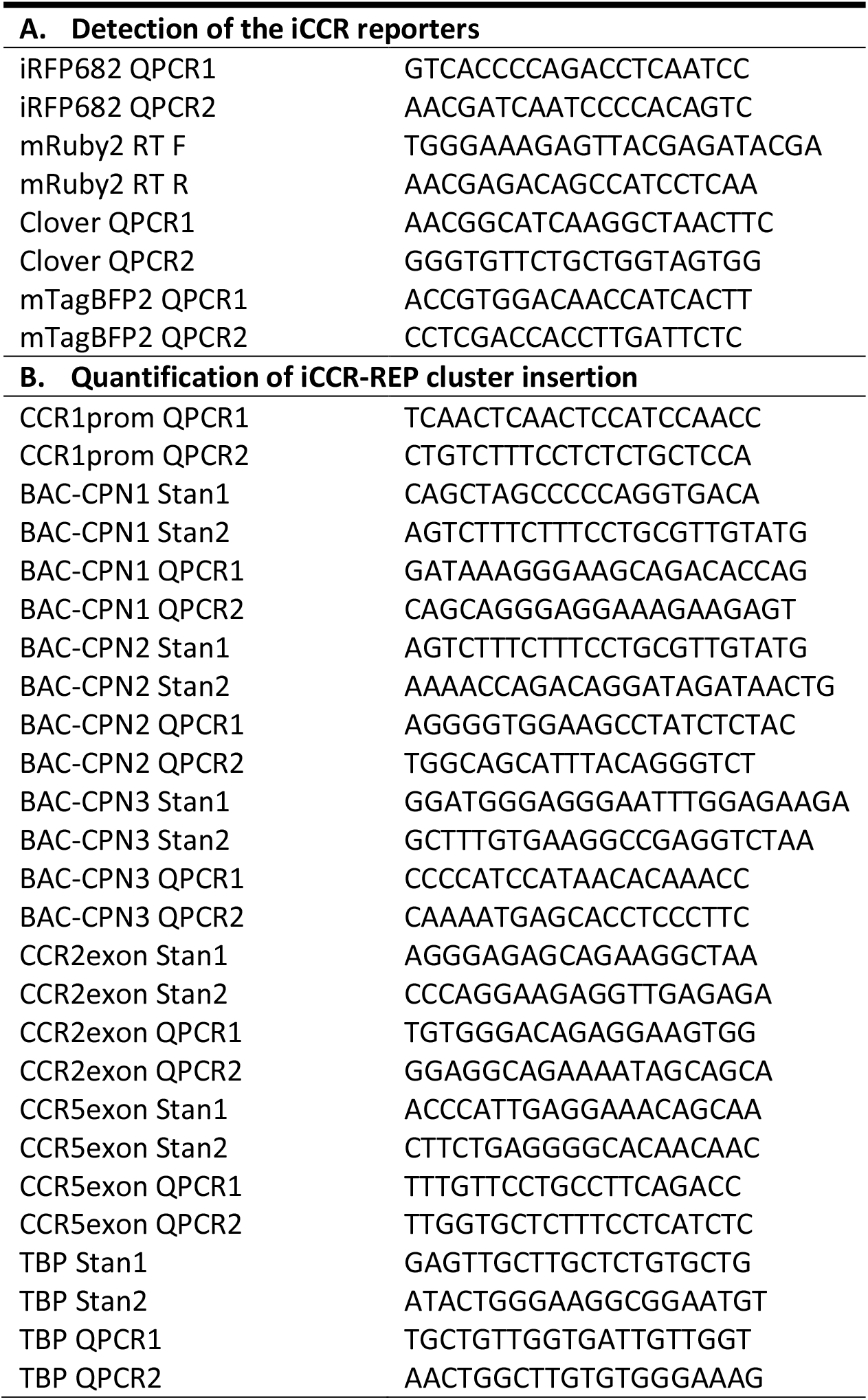
Primers used in the study

**Table S2.** Antibodies used in the study

| Antibody | Clone | Source | Working dilution |
| --- | --- | --- | --- |
| Anti-mouse CD45 | 30-F11 | eBioscience | 1/100 |
| Anti-mouse CD11b | M1/70 | eBioscience | 1/100 |
| Anti-mouse SiglecF | E50-2440 | BD Bioscience | 1/100 |
| Anti-mouse F4/80 | BM8 | eBioscience | 1/100 |
| Anti-mouse CD64 | X54-5/7.1 | BD Bioscience | 1/100 |
| Anti-mouse Ly6C | HK1.4 | BioLegend | 1/100 |
| Anti-mouse CD11c | HL3 | BD Bioscience | 1/100 |
| Anti-mouse MHCII | M5/114.15.2 | BioLegend | 1/100 |
| Anti-mouse Ly6G | 1A8 | BD Bioscience | 1/100 |
| Anti-mouse CD19 | eBio1D3 (1D3) | eBioscience | 1/100 |
| Anti-mouse CCR2 | SA203G11 | BioLegend | 1/50 |
| Anti-mouse CCR3 | J073E5 | BioLegend | 1/50 |
| Anti-mouse CCR5 | HM-CCR5 (7A4) | eBioscience | 1/50 |

